## Supplemental Material for "3-O-methyltolcapone and Its Lipophilic Analogues are Potent Inhibitors of Transthyretin Amyloidogenesis with High Permeability and Low Toxicity"

^h^Tes Pharma S.r.l., Via P. Togliatti 20, Corciano, 06073, Perugia, Italy.

^i^Department of Food and Drug, University of Parma, 43124 Parma, Italy

^l^Department of Agricultural, Food and Environmental Sciences, Università Politecnica delle Marche, Via Brecce Bianche, 60131 Ancona, Italy

^#^current address: Dept. of Pathobiology, Faculty of Science, Mahidol University, Rama VI Road, Ratchathewi, Bangkok 10400 THAILAND

Corresponding Author

### **Table S1**. Summary table of molecular docking performed on five analogs of 3-O-methyltolcapone and the tetrameric structure of apoTTR (PDB ID: 4D7B) and LogP values of the 3-O-methyltolcapone analogs calculated using the software Spartan (Wavefunction, Inc. Spartan’16 (Irvine, CA), Wavefunction, Inc.))

| Compound* | Number of runs | Total number of clusters | FF of the top cluster binding HBPs (Kcal/mol) | ΔG of the top cluster binding HBPs (Kcal/mol) | LogP |
| --- | --- | --- | --- | --- | --- |
| Tolcapone# |  |  |  |  | 2.99 |
| 3-O-methyltolcapone | 3 | 32 | -2702 | -7.18 | 3.25 |
| **1** | 3 | 35 | -2705 | -7.70 | 3.74 |
| **2** | 3 | 34 | -2704 | -7.65 | 3.74 |
| **3** | 3 | 36 | -2668 | -7.18 | 3.74 |
| **4** | 3 | 30 | -2694 | -7.12 | 4.22 |
| **5** | no binding | no binding | no binding | no binding | 4.22 |

* Three docking runs were performed for each compound. The resulting clusters were listed based on the lowest FF score. The cluster binding in the HBPs and with the lowest values was used for the evaluation of the protein-ligand interaction. The table reports the best values from the three runs with the ligand binding the HBPs.

\# docking was not performed on this compound.

### **Scheme S1**. Attempt of synthesis of compound 2 through strategy A. Reaction conditions: i) magnesium turnings, 2,4-dimethylbromobenzene, dry THF, rt, 1.5 h; ii) tBuONa, cyclohexanone, toluene, reflux, 16 h; iii) HCOO-NH4+, Pd/C, MeOH, reflux, 0.5 H; iv) glacial AcOH, 65% HNO3, rt, 0.5 h.

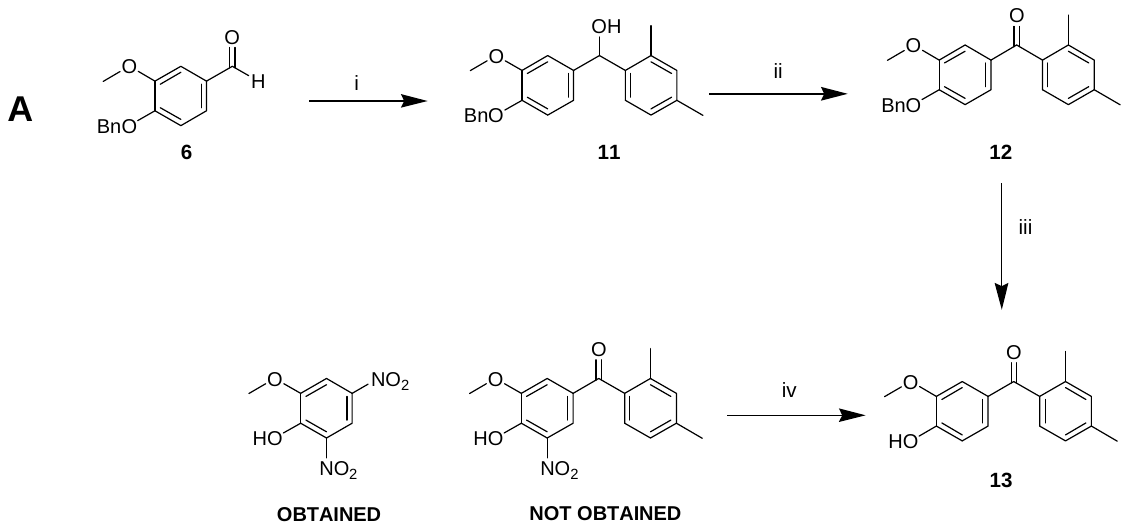

**(4-(benzyloxy)-3-methoxyphenyl)-1-(2,4-dimethylphenyl)methanol (11).** In a flame-dried three-necked round-bottomed flask, magnesium turnings (0.30 g, 12.52 mmol) were suspended in dry THF (3 mL) under Ar atmosphere. Using a dropping funnel, a solution in dry THF (2.3 mL) of 1-bromo-2,4-dimethylbenzene (2.31 g, 12.52 mmol) was slowly added in 1 hour. Then, a solution of **6** (2 g, 8.26 mmol) in dry THF (1.7 mL) was slowly dripped in 30 min into the stirred reaction mixture. The reaction progression was monitored by TLC (eluent: hexane/ethyl acetate 4/1). After 1 hour, the reaction was quenched by addition of saturated NH_4_Cl solution (5 mL), extracted with Et_2_O (3x10 mL) and then the organic phase was washed with brine (3x10 mL). The organic phase was dried over anhydrous Na_2_SO_4_, evaporated under reduced pressure and the residue was crystallized from Et_2_O/petroleum ether to afford the product as white solid (1.6 g, 56%) of m.p. 59-60 °C. ^1^H NMR (DMSO-d_6_,400 MHz) δ 7.45 – 7.27 (m, 6H, ArH); 6.98 (brd, 1H, ArH); 6.96 – 6.88 (m, 3H, ArH); 6.67 (dd, *J* = 8.3, 2.0 Hz, 1H, ArH); 5.73 (d, *J* = 4.3 Hz, 1H, OH); 5.58 (d, *J* = 4.4 Hz, 1H, CH); 5.03 (s, 2H, CH_2_); 3.72 (s, 3H, OCH_3_); 2.24 (s, 3H, CH_3_); 2.17 (s, 3H, CH_3_). ^13^C NMR (DMSO-d_6_,100 MHz) δ 149.2 (Ar), 147.0 (Ar), 140.8 (Ar), 138.2 (Ar), 137.7 (Ar), 135.9 (Ar), 135.0 (Ar), 131.1 (Ar), 128.8 (Ar), 128.2 (Ar), 128.2 (Ar), 126.9 (Ar), 126.6 (Ar), 119.4 (Ar), 113.7 (Ar), 111.7 (Ar), 71.5 (CH), 70.4 (CH_2_), 56.0 (OCH_3_), 21.0 (CH_3_), 19.5 (CH_3_). MS (ESI, m/z): 371.13 [M + Na]^+^.

**(4-(benzyloxy)-3-methoxyphenyl)-1-(2,4-dimethylphenyl)methanone (12).** t-BuONa (0.22 g, 2.32 mmol) and cyclohexanone (0.94 mL, 9.09 mmol) were added to a solution of **11** (0.49 g, 1.40 mmol) in toluene (2 mL). The reaction progression was monitored by TLC (eluent: hexane/ethyl acetate 7/3). The solution was stirred at reflux for 16 h. The solution was cooled at 50 °C and then water (2 mL) was added. The organic phase was separated from water and the aqueous phase was extracted with ethyl acetate (3x3 mL). The combined organic phases were collected and washed with water (10 mL) and brine (10 mL). Subsequently, the solvent was evaporated under reduced pressure obtaining an oily residue that was crystallized with ethanol 96% to afford **12** as white powder (0.27 g, 57%) of m.p. 100-101 °C. ^1^H NMR (CDCl_3_,400 MHz): δ 7.58 (d, *J* = 2.0 Hz, 1H, ArH); 7.47 – 7.31 (m, 5H, ArH); 7.24 – 7.17 (m, 2H, ArH); 7.11 (brs, 1H, ArH); 7.05 (d, 1H, ArH); 6.86 (d, *J* = 8.4 Hz, 1H, ArH); 5.25 (s, 2H, CH_2_); 3.97 (s, 3H, OCH_3_); 2.39 (s, 3H, CH_3_); 2.31 (s, 3H, CH_3_). ^13^C NMR (CDCl_3_, 100 MHz) δ 197.3 (C=O), 158.3 (Ar), 152.5 (Ar), 149.5 (Ar), 140.1 (Ar), 136.6 (Ar), 136.3 (Ar), 136.0 (Ar), 131.7 (Ar), 131.2 (Ar), 128.5 (Ar), 128.1 (Ar), 127.2 (Ar), 125.7 (Ar), 125.7 (Ar), 111.9 (Ar), 111.7 (Ar), 70.8 (CH_2_), 56.1 (OCH_3_), 21.4 (CH_3_), 19.9 (CH_3_). MS (ESI, m/z): 369.11 [M + Na]^+^.

**(4-hydroxy-3-methoxyphenyl)-1-(2,4-dimethylphenyl)methanone (13).** To a solution in methanol (2.4 mL) of **12** (0.27 g, 0.79 mmol) and ammonium formate (0.20 g, 3.16 mmol), Pd/C 10% (catalytic amount) suspended in methanol (1.0 mL) was added. The resulting mixture was stirred at reflux for 1 hour. The reaction progression was monitored by TLC (eluent: hexane/ethyl acetate 7/3). Subsequently, it was cooled in an ice/water bath and water (1 mL) and HCl 2M (0.2 mL) were slowly added up to slightly acidic pH. After addition of DCM (3mL), the mixture was filtered through a celite pad. The organic phase was separated and the aqueous one was extracted with DCM (3x 3mL). The combined organic phases were washed with water (10 mL) and brine (10 mL) and dried with anhydrous Na_2_SO_4_. The solvent was evaporated under reduced pressure and the product **13** was obtained by crystallization from DCM/petroleum ether as yellow crystals (0.19 g, 94%) of m.p. 116-117 °C. ^1^H NMR (400 MHz, CDCl_3_): 7.58 (d, *J* = 1.9 Hz, 1H, H_a_); 7.24 – 7.20 (m, 2H, H_c_, H_d_); 7.12 (brs, 1H, H_f_); 7.06 (brd, 1H, H_e_); 6.91 (d, *J* = 8.2 Hz, 1H, H_b_); 6.09 (s, 1H, OH); 3.99 (s, 3H, OCH_3_); 2.40 (s, 3H, CH_3_); 2.31 (s, 3H, CH_3_). ^13^C NMR (100 MHz, CDCl_3_): δ(ppm) = 197.4 (C=O), 150.4 (Ar), 146.6 (Ar), 140.0 (Ar), 136.5 (Ar), 136.1 (Ar), 131.7 (C-H_f_), 130.7 (Ar), 128.4 (Ar), 126.7 (Ar), 125.7 (C-H_e_), 113.6 (C-H_b_), 110.9 (C-H_a_), 56.1 (OCH_3_), 21.4 (CH_3_), 19.8 (CH_3_). MS (ESI, m/z): 257.10 [M + Na]^+^.

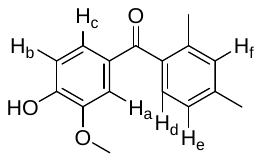

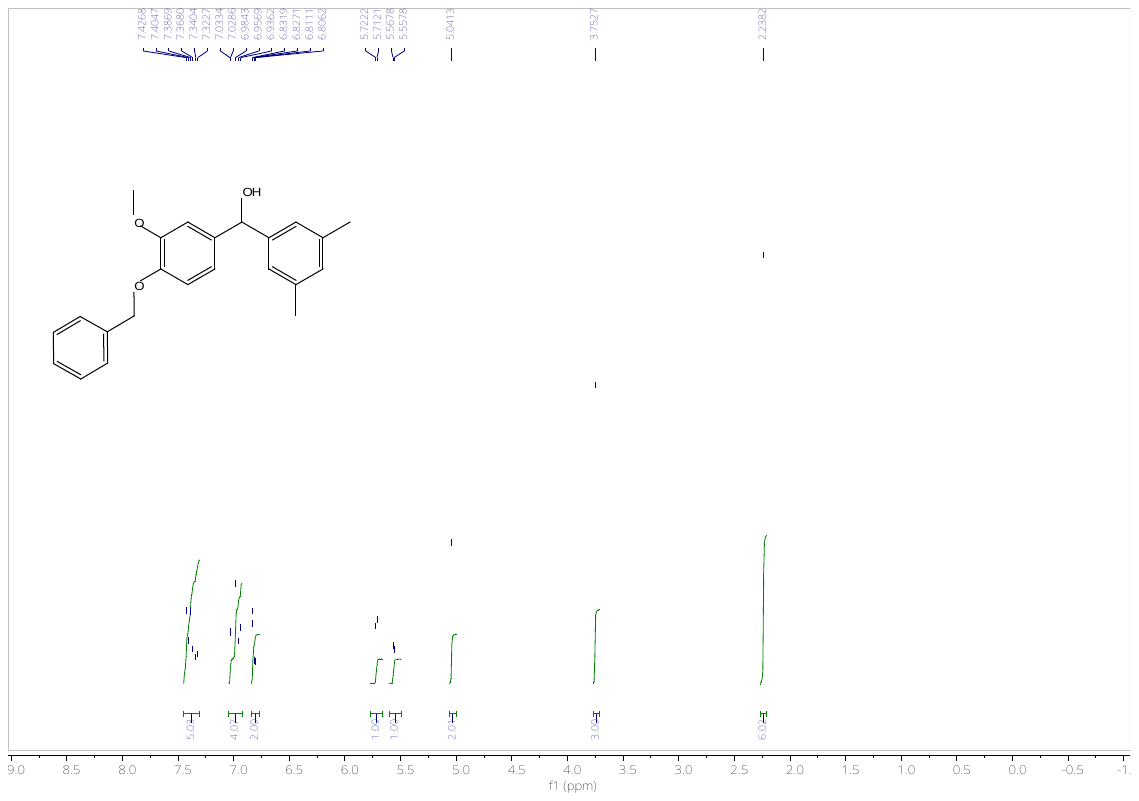

**^1^H NMR** spectrum of **7** (400 MHz, DMSO-d_6_, 298K)

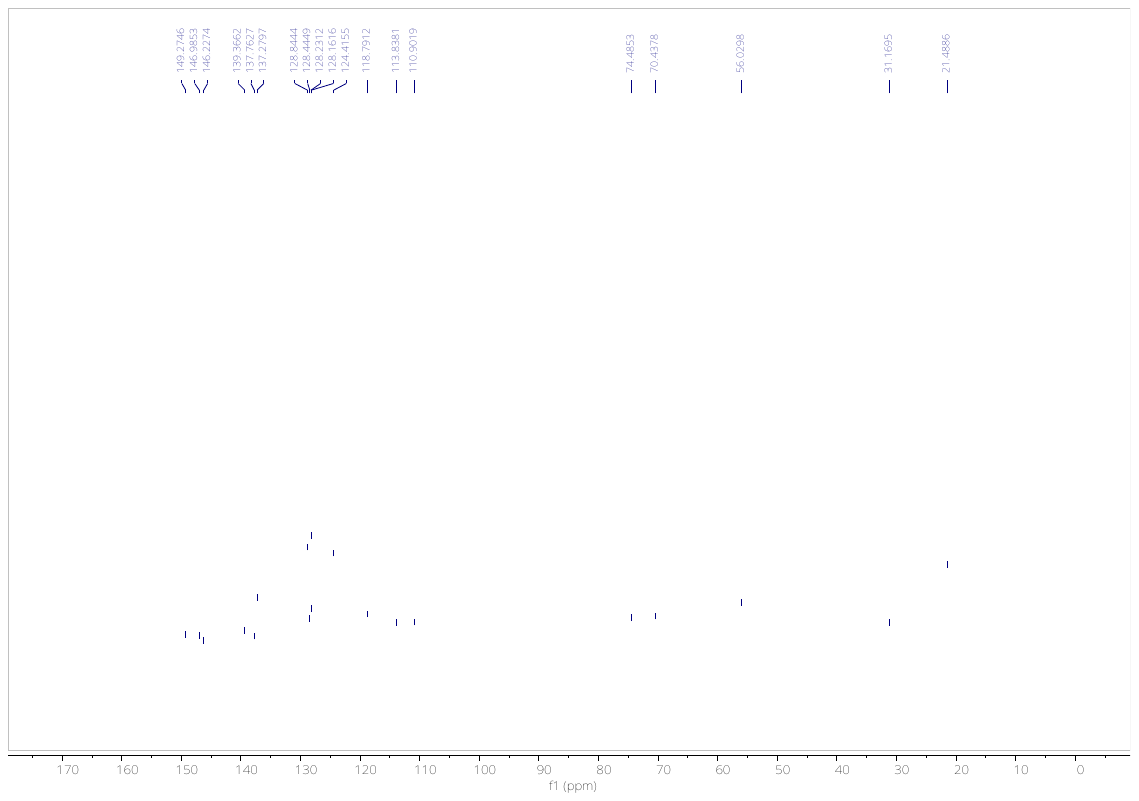

**^13^C NMR** spectrum of **7** (100 MHz, DMSO-d_6_, 298K)

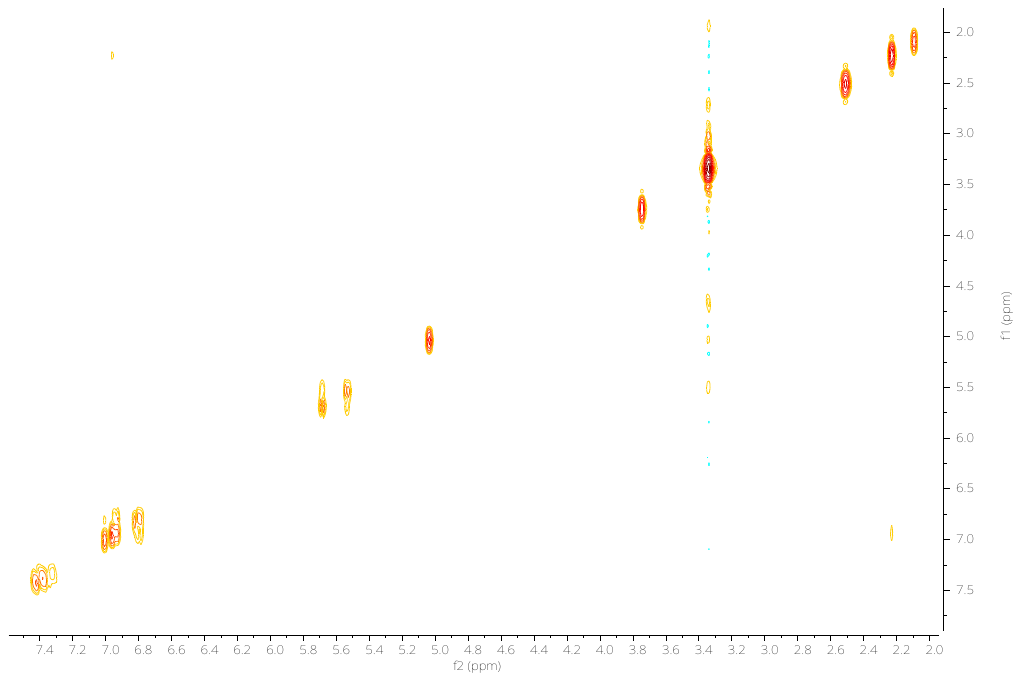

**^1^H-^1^H COSY NMR** spectrum of **7** (400 MHz, DMSO-d_6_, 298K)

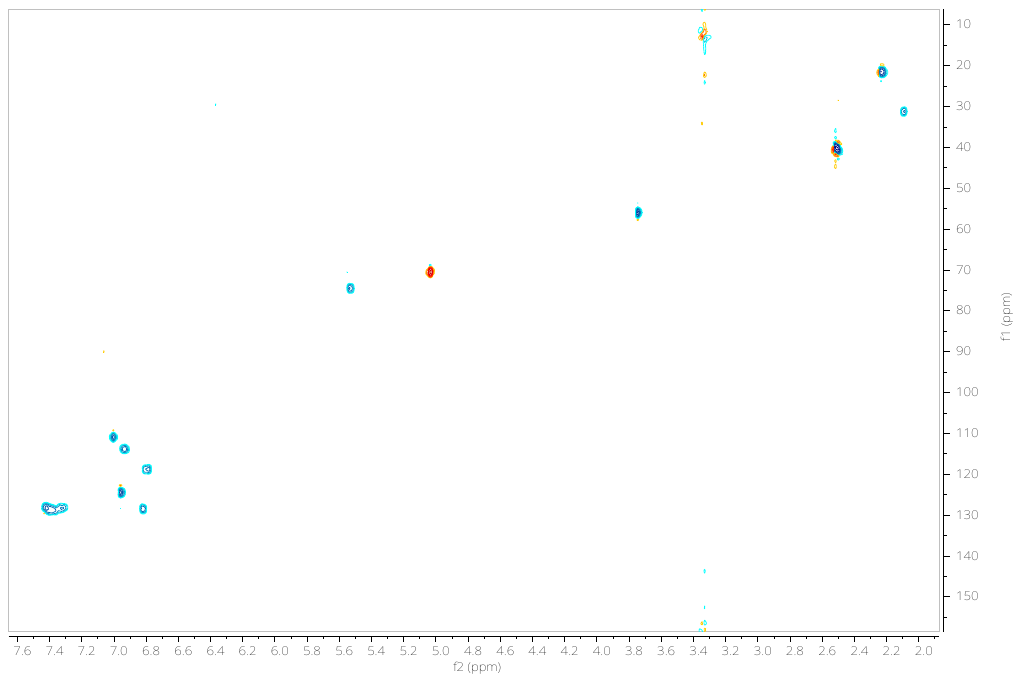

**^1^H-^13^C HSQC NMR** spectrum of **7** (DMSO-d_6_, 298K)

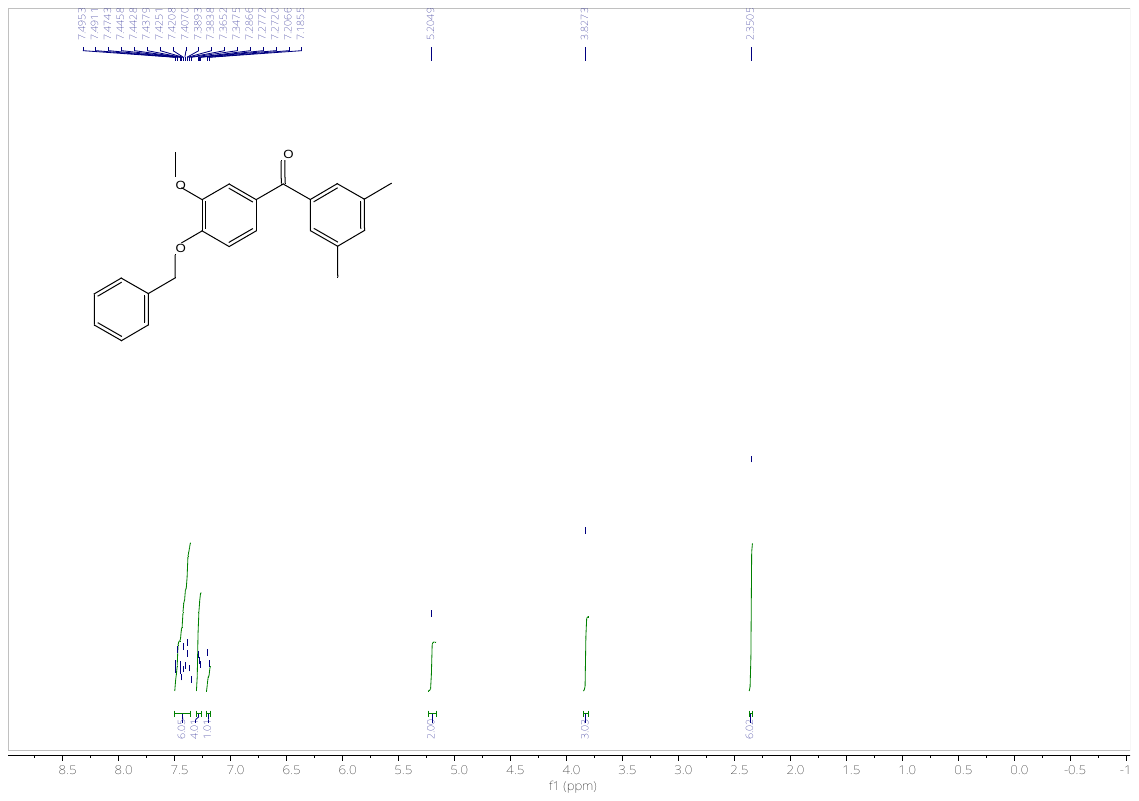

**^1^H NMR** spectrum of **8** (400 MHz, DMSO-d_6_, 298K)

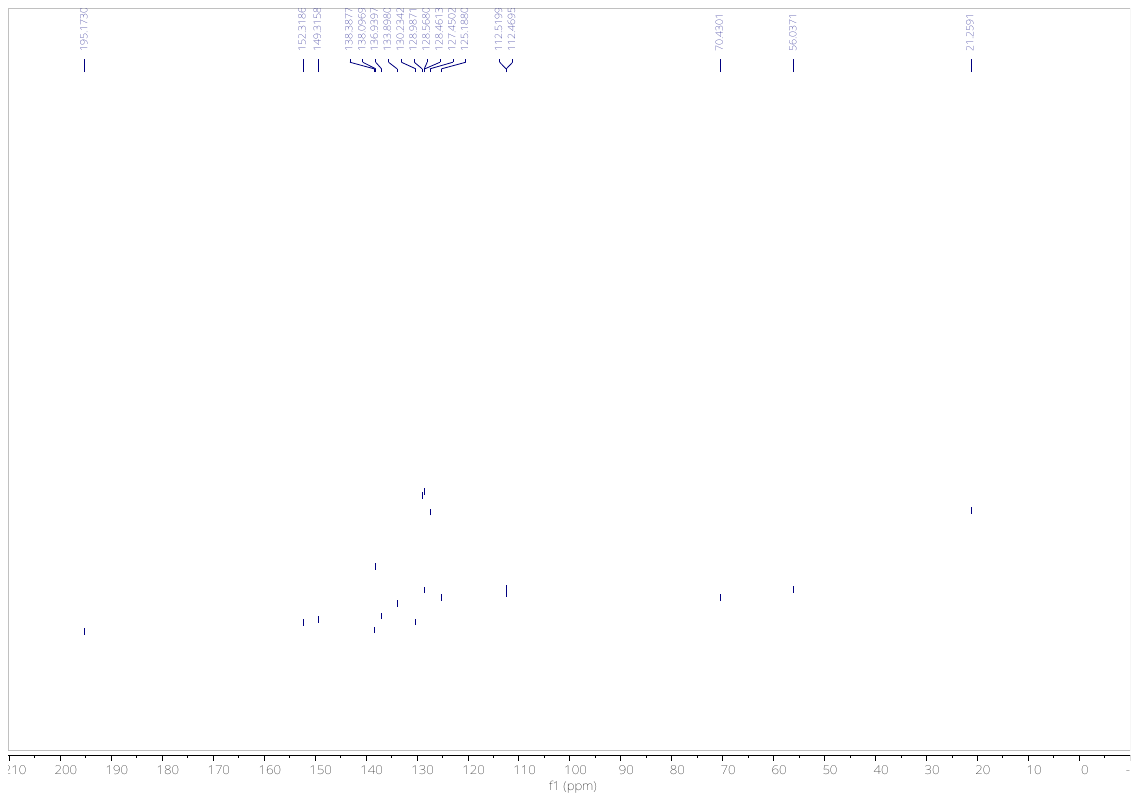

**^13^C NMR** spectrum of **8** (100 MHz, DMSO-d_6_, 298K)

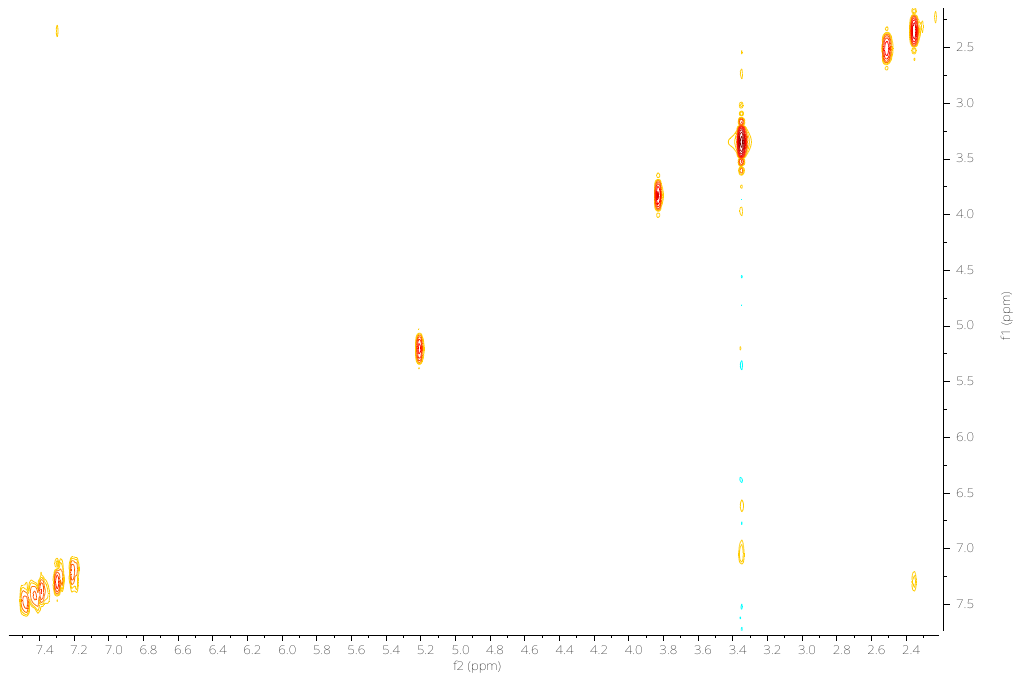

**^1^H-^1^H COSY NMR** spectrum of **8** (400 MHz, DMSO-d_6_, 298K)

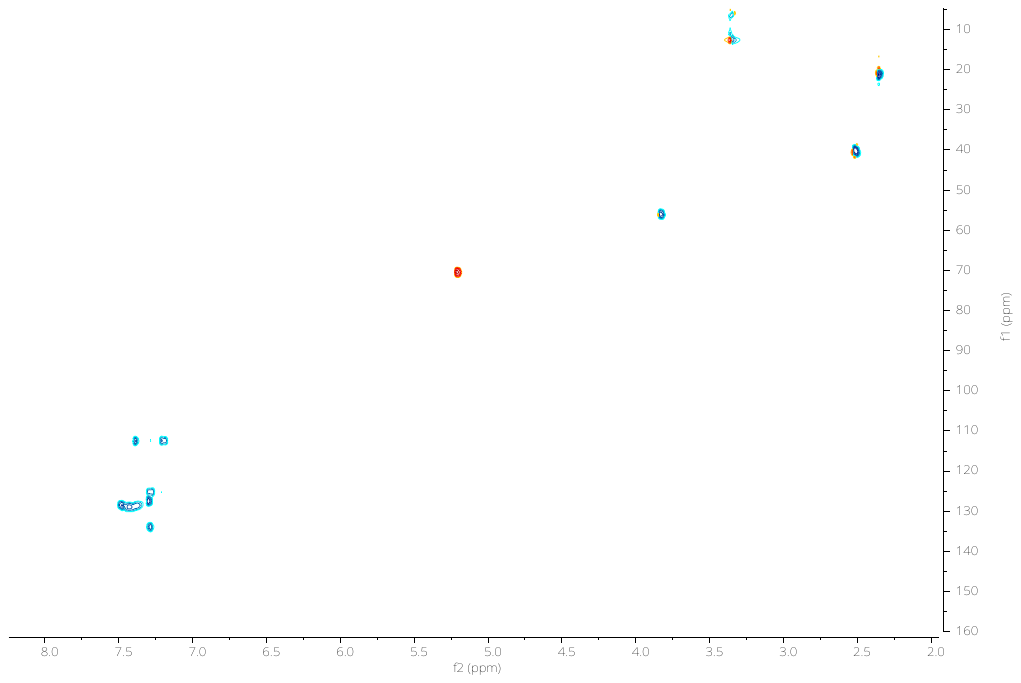

**^1^H-^13^C HSQC NMR** spectrum of **8** (DMSO-d_6_, 298K)

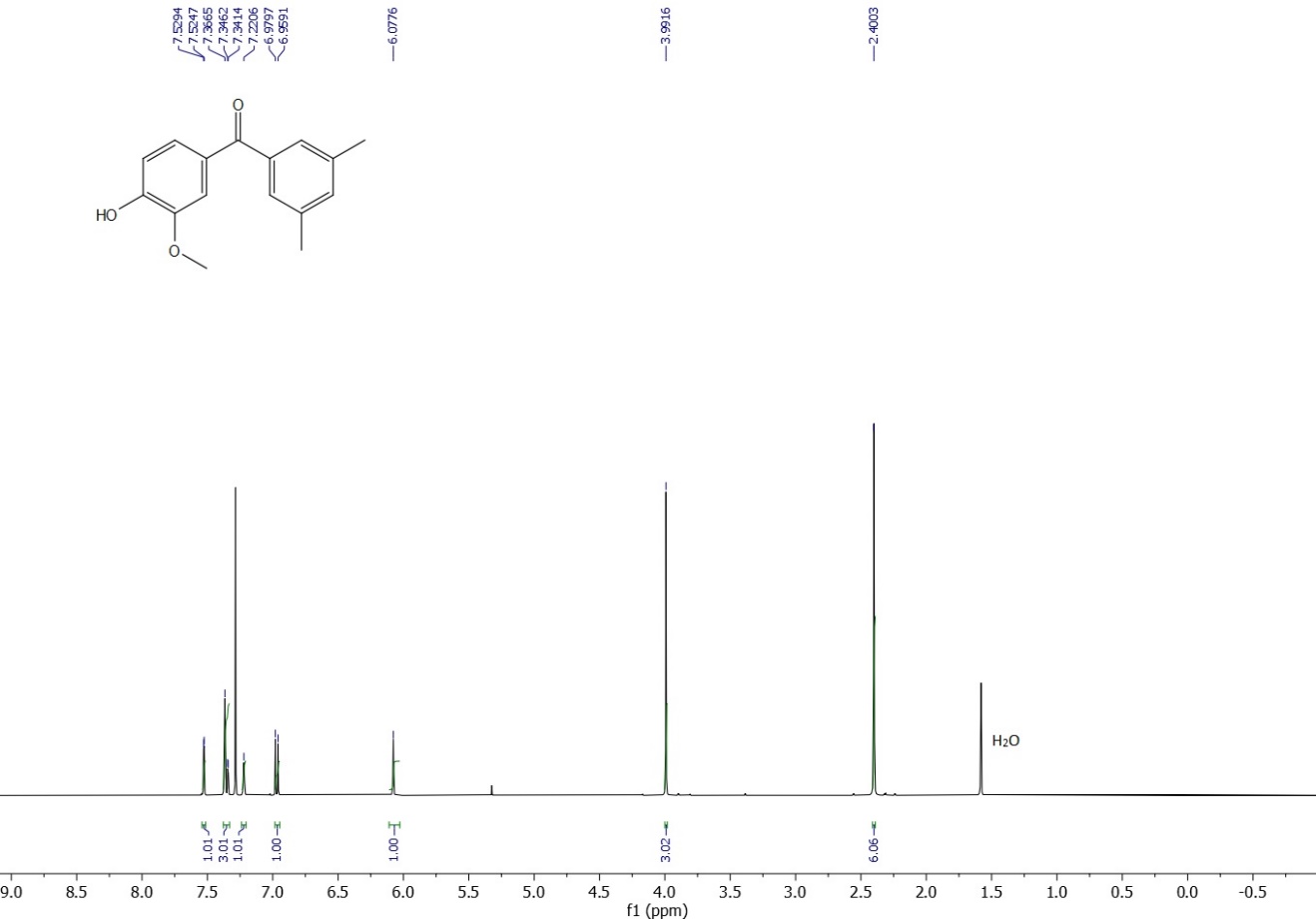

**^1^H NMR** spectrum of **9** (400 MHz, CDCl_3_, 298K)

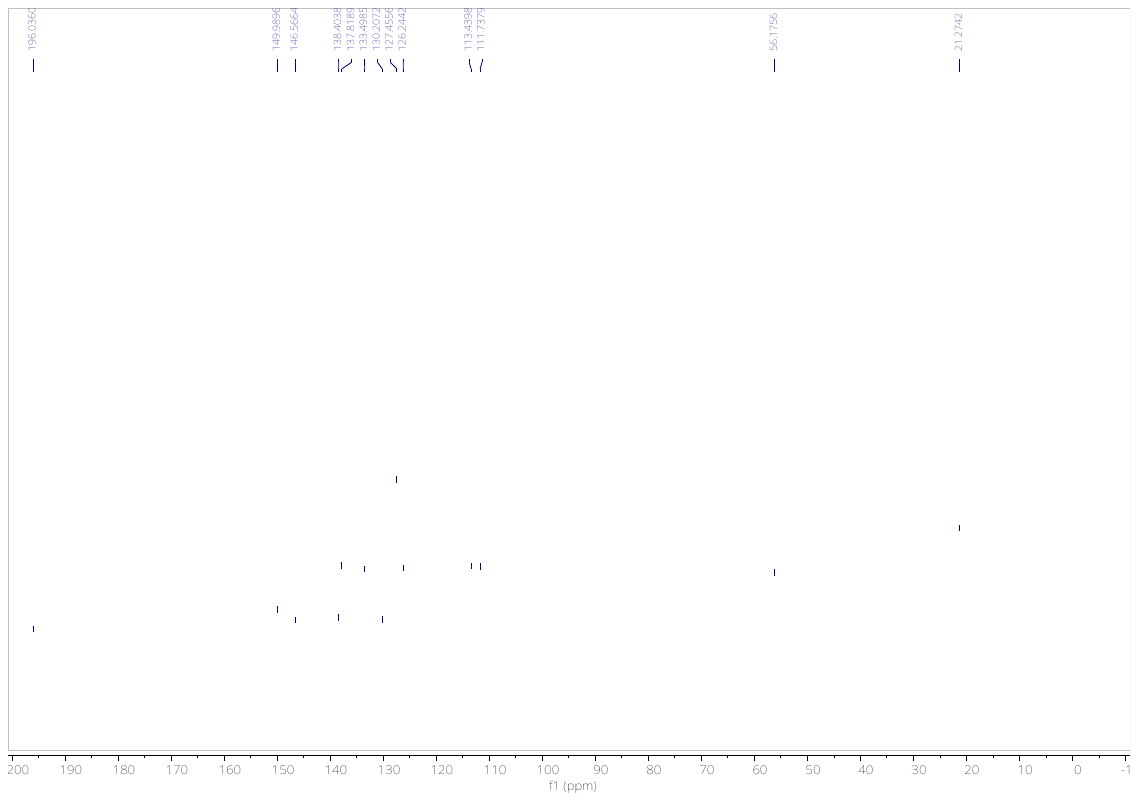

**^13^C NMR** spectrum of **9** (100 MHz, CDCl_3_, 298K)

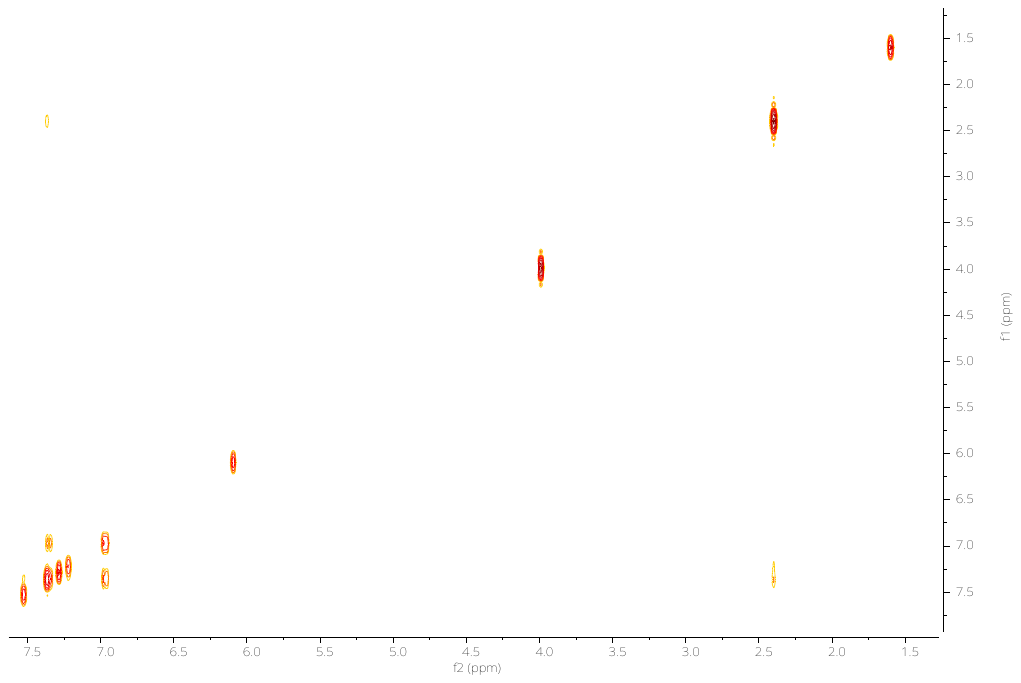

**^1^H-^1^H COSY NMR** spectrum of **9** (400 MHz, CDCl_3_, 298K)

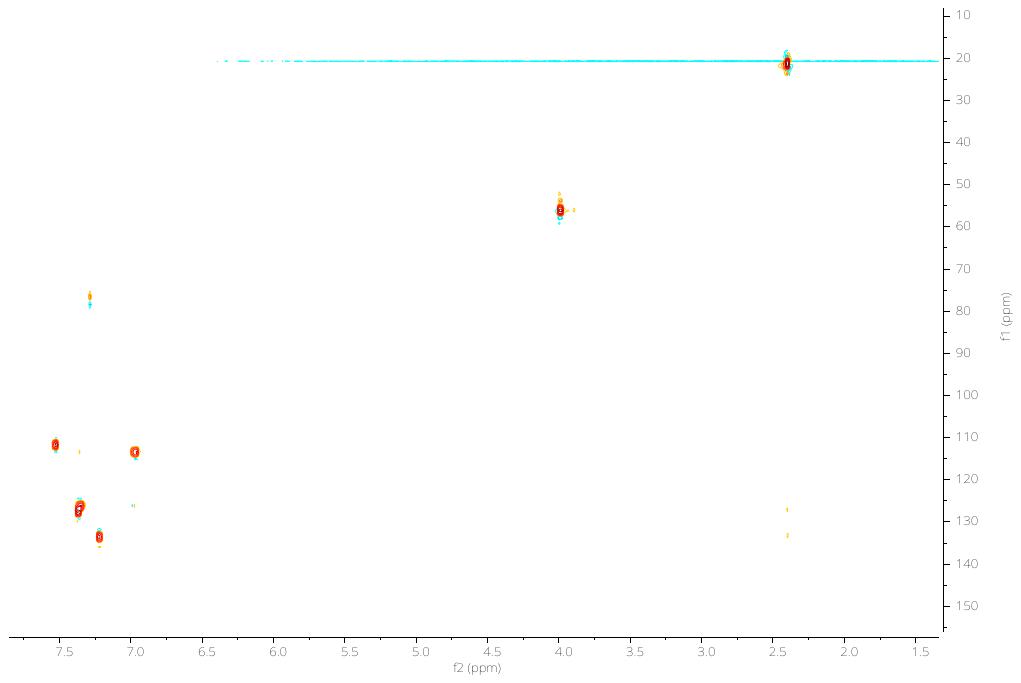

**^1^H-^13^C HSQC NMR** spectrum of **9** (CDCl_3_, 298K)

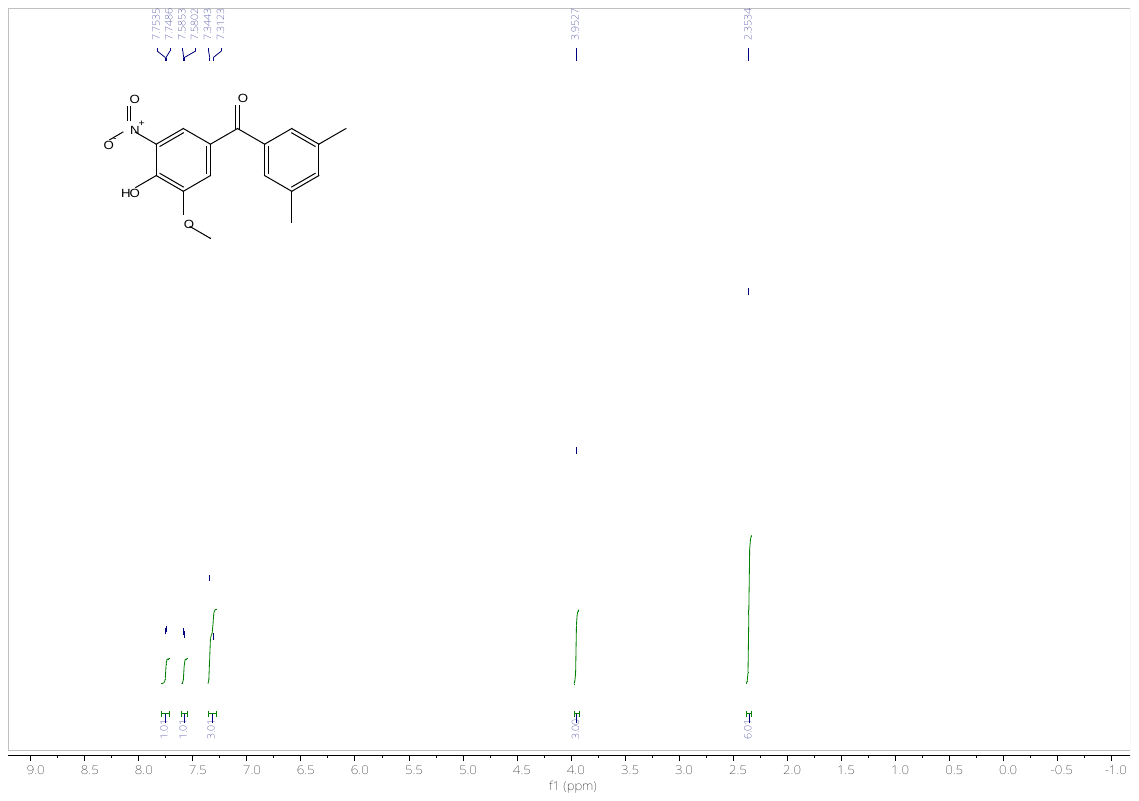

**^1^H NMR** spectrum of **1** (400 MHz, DMSO-d_6_, 298K)

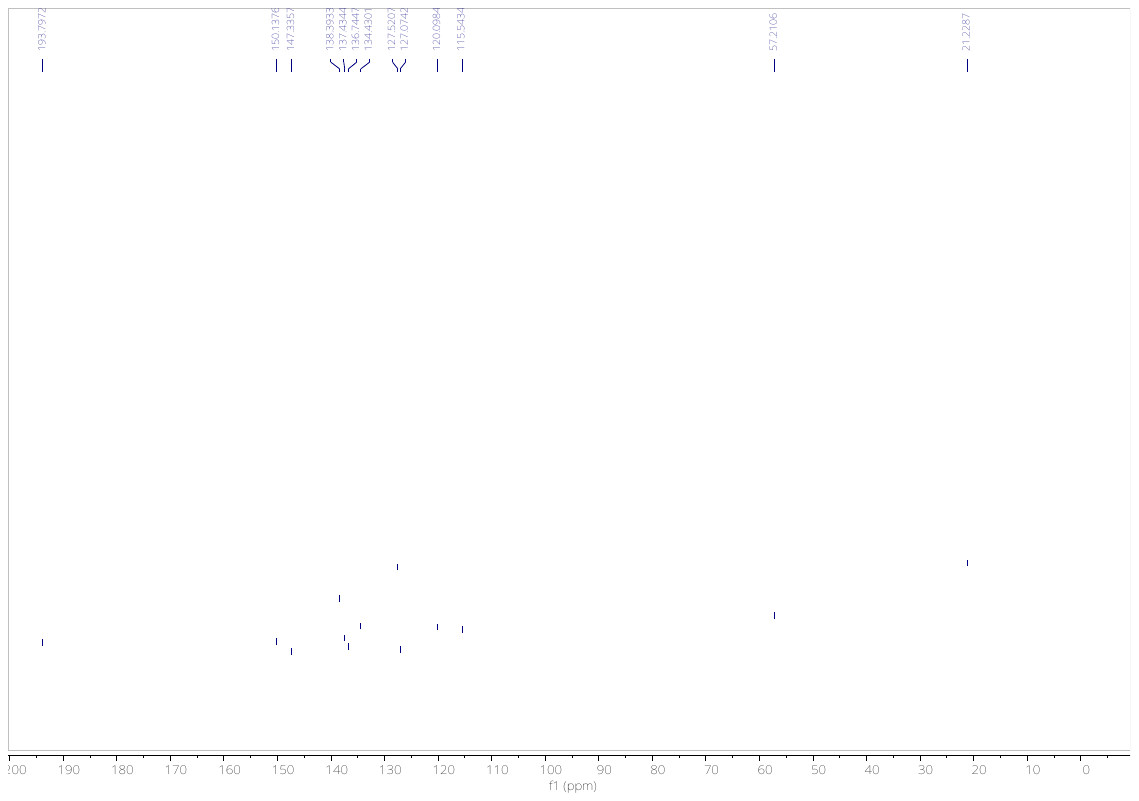

**^13^C NMR** spectrum of **1** (100 MHz, DMSO-d_6_, 298K)

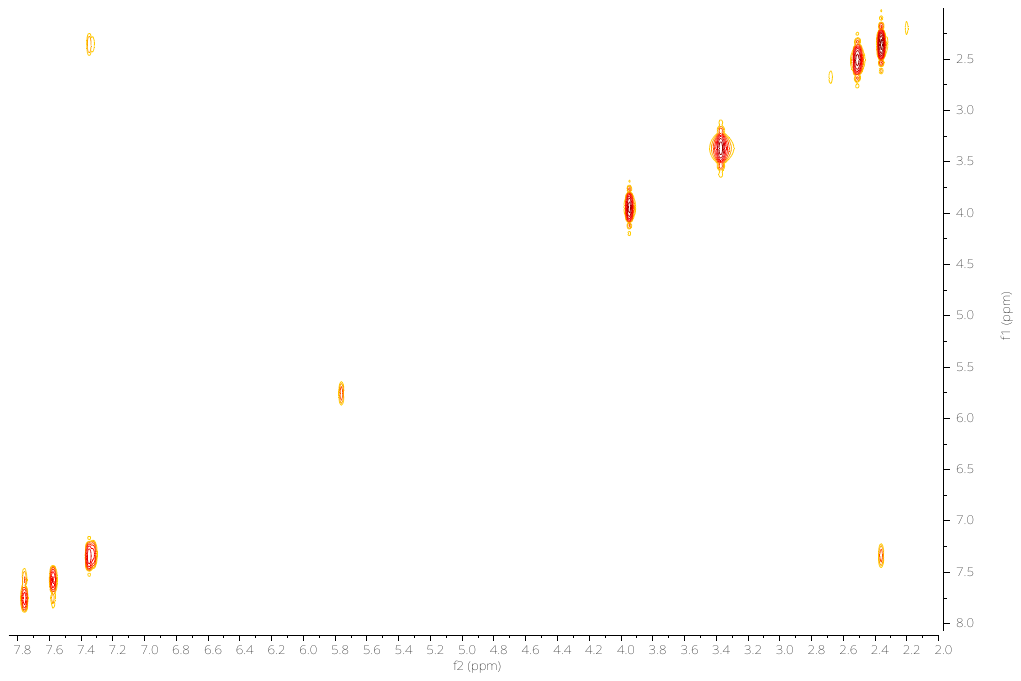

**^1^H-^1^H COSY NMR** spectrum of **1** (400 MHz, DMSO-d_6_, 298K)

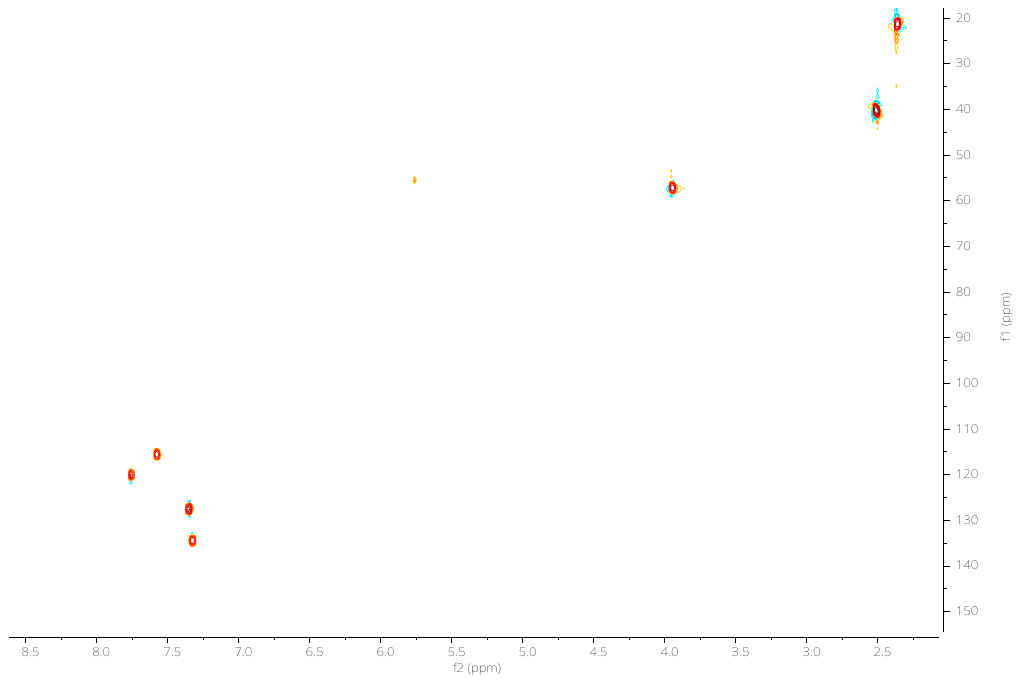

**^1^H-^13^C HSQC NMR** spectrum of **1** (DMSO-d_6_, 298K)

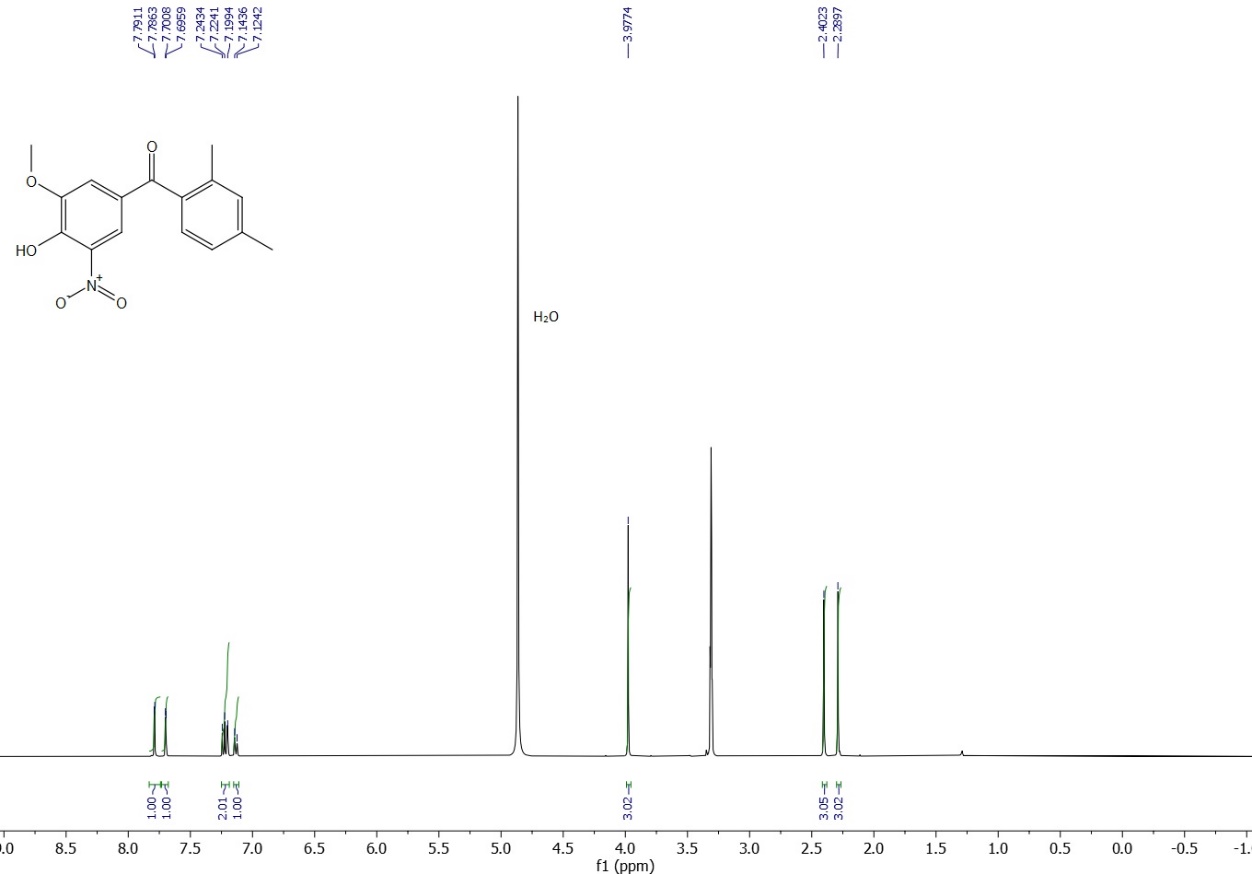

**^1^H NMR** spectrum of **2** (400 MHz, CD_3_OD, 298K)

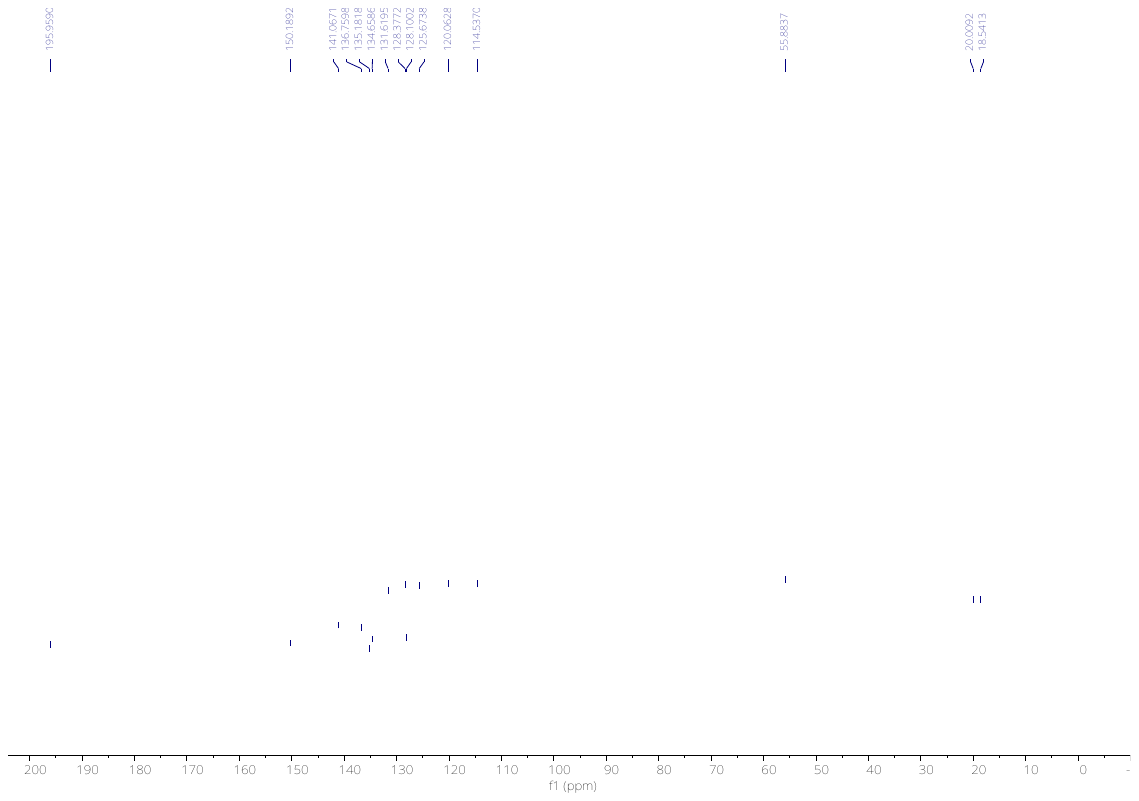

**^13^C NMR** spectrum of **2** (100 MHz, CD_3_OD, 298K)

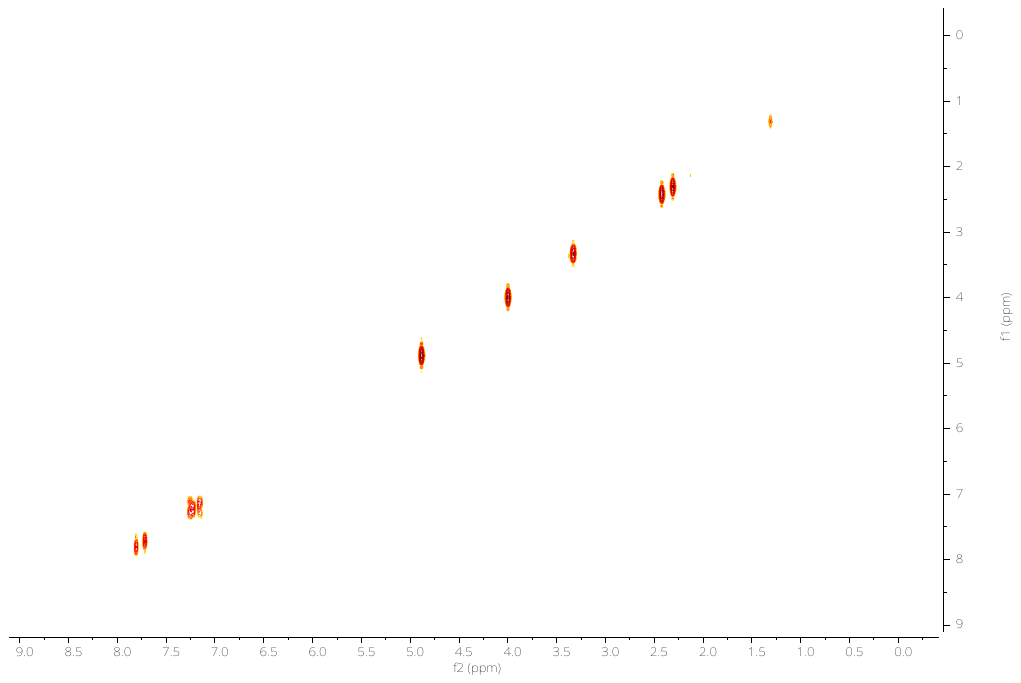

**^1^H-^1^H COSY NMR** spectrum of **2** (400 MHz, CD_3_OD, 298K)

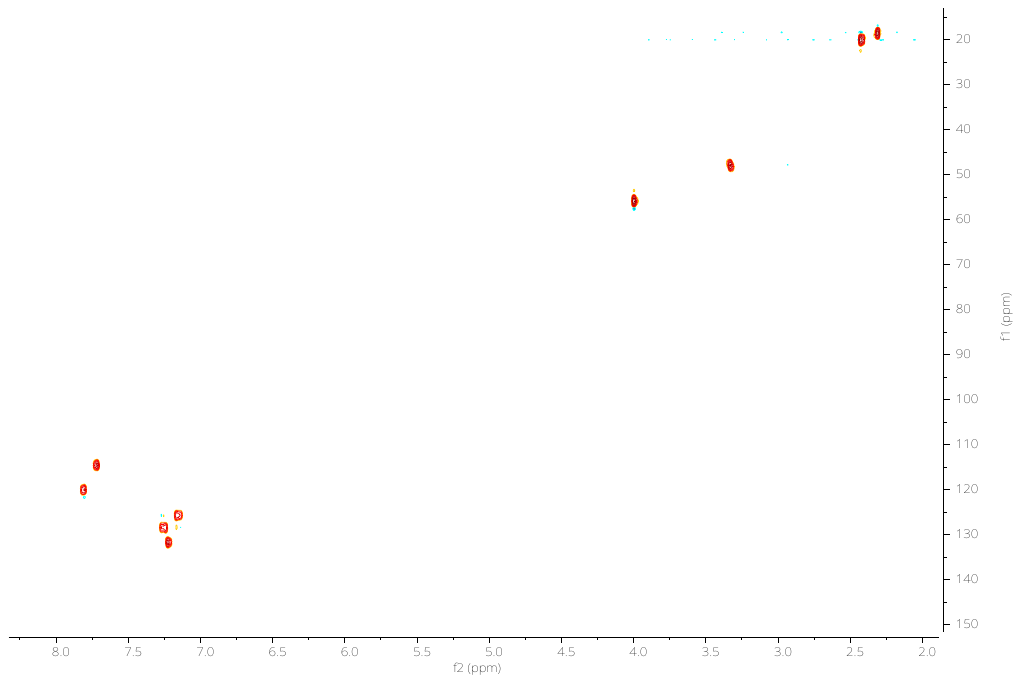

**^1^H-^13^C HSQC NMR** spectrum of **2** (CD_3_OD, 298K)

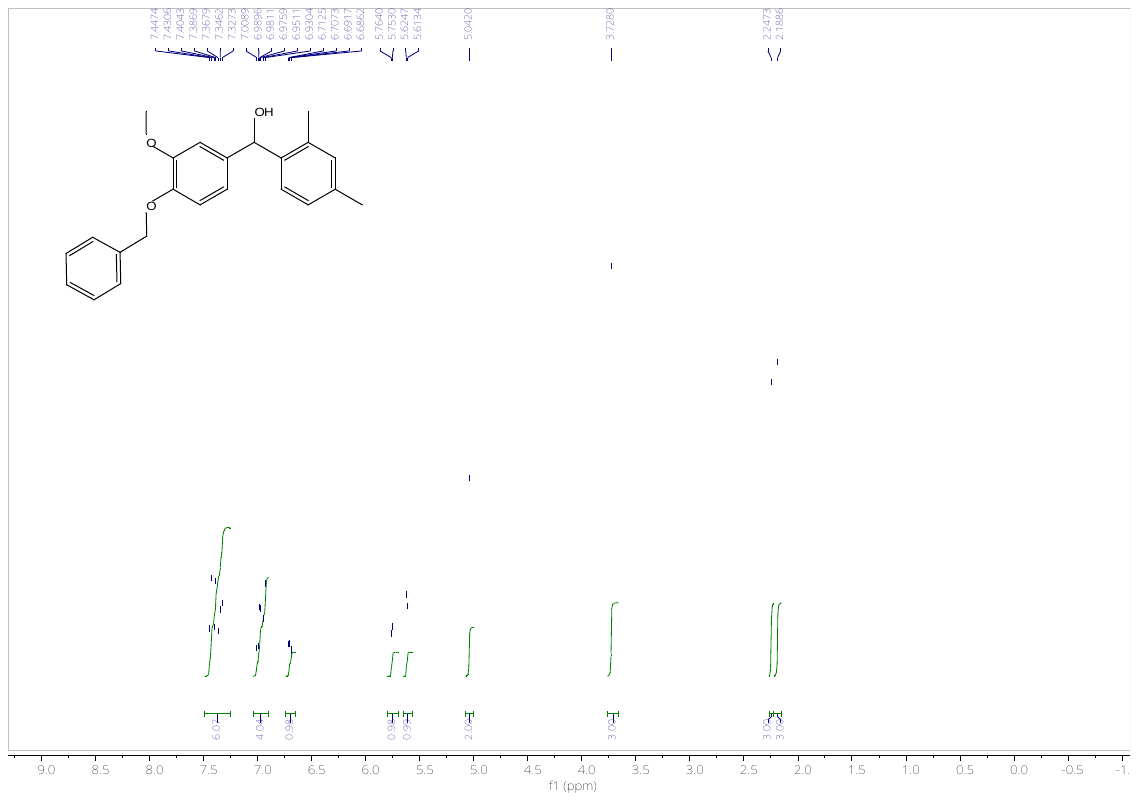

**^1^H NMR** spectrum of **11** (400 MHz, DMSO-d_6_, 298K)

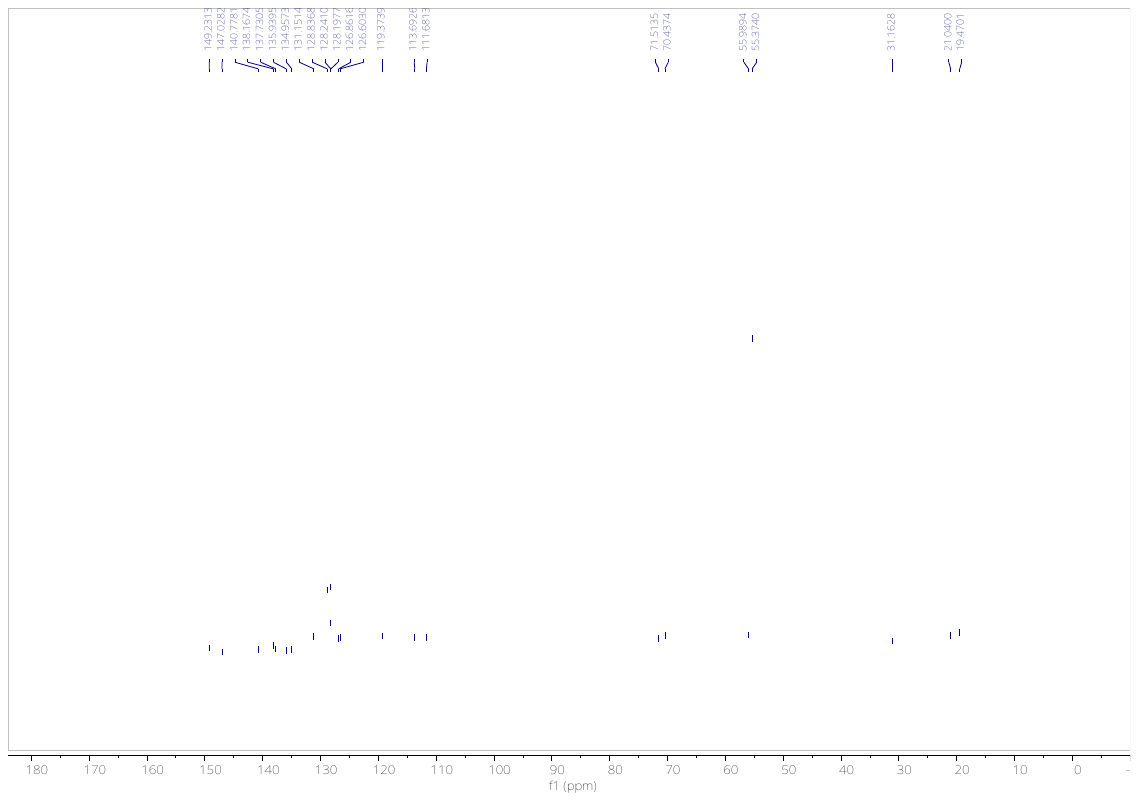

**^13^C NMR** spectrum of **11** (100 MHz, DMSO-d_6_, 298K)

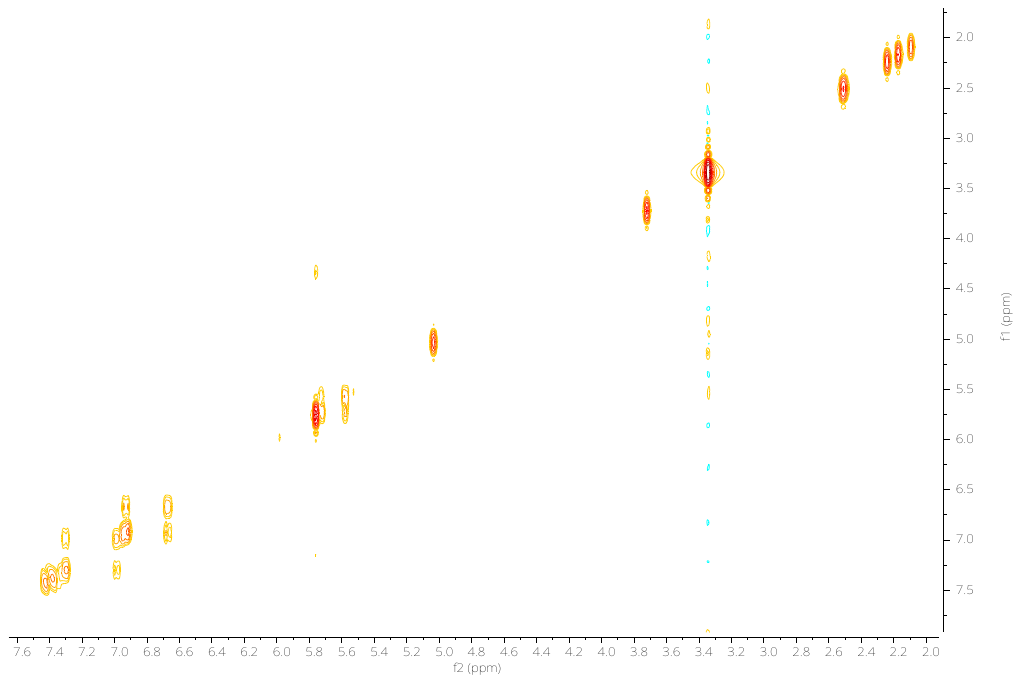

**^1^H-^1^H COSY NMR** spectrum of **11** (400 MHz, DMSO-d_6_, 298K)

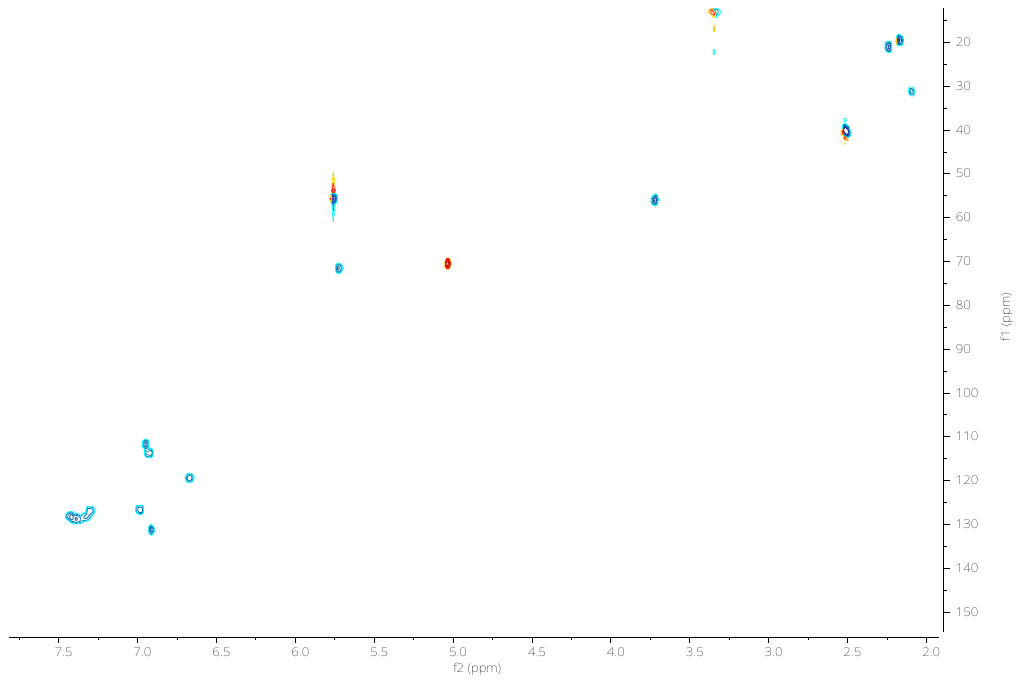

**^1^H-^13^C HSQC NMR** spectrum of **11** (DMSO-d_6_, 298K)

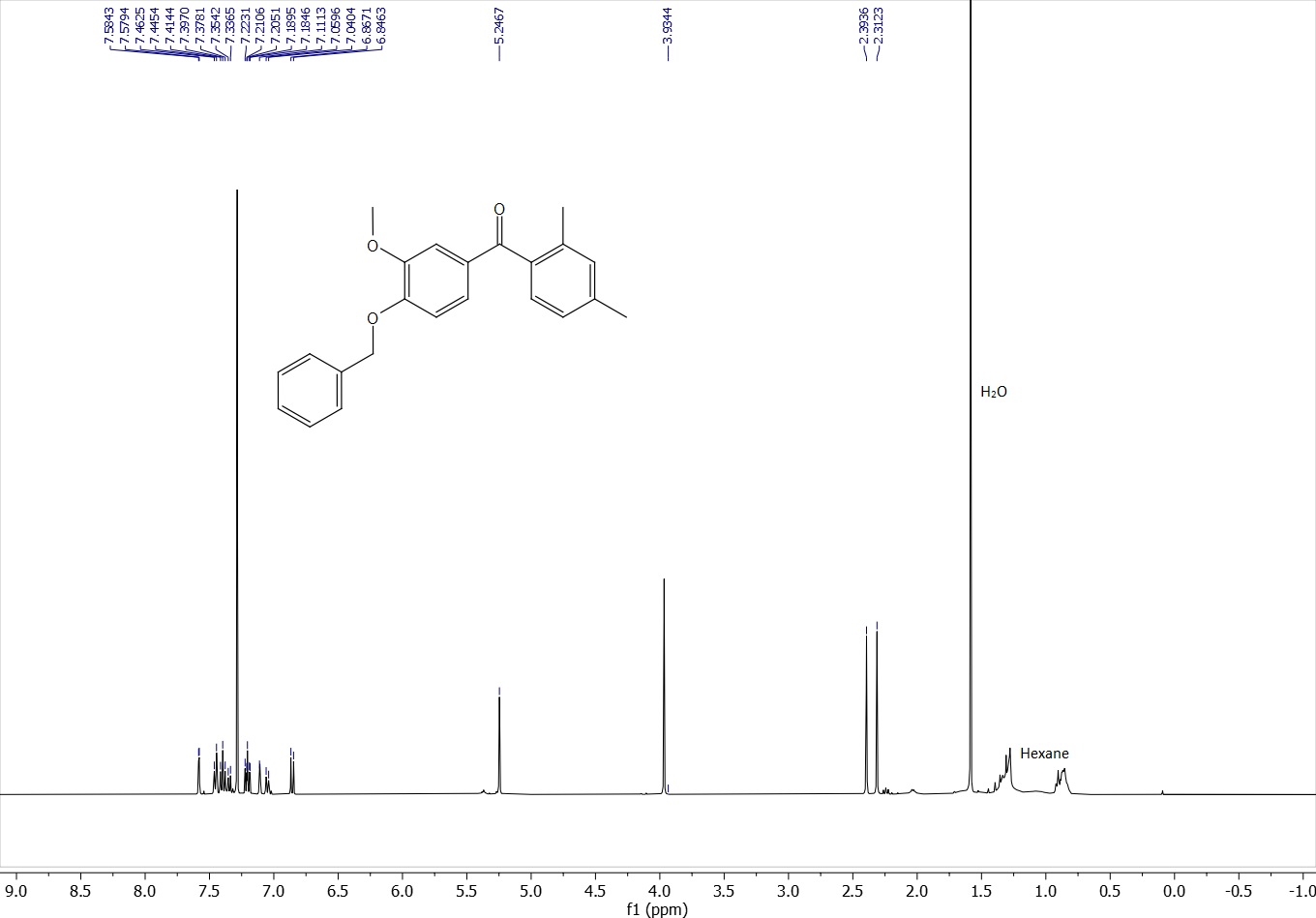

**^1^H NMR** spectrum of **12** (400 MHz, CDCl_3_, 298K)

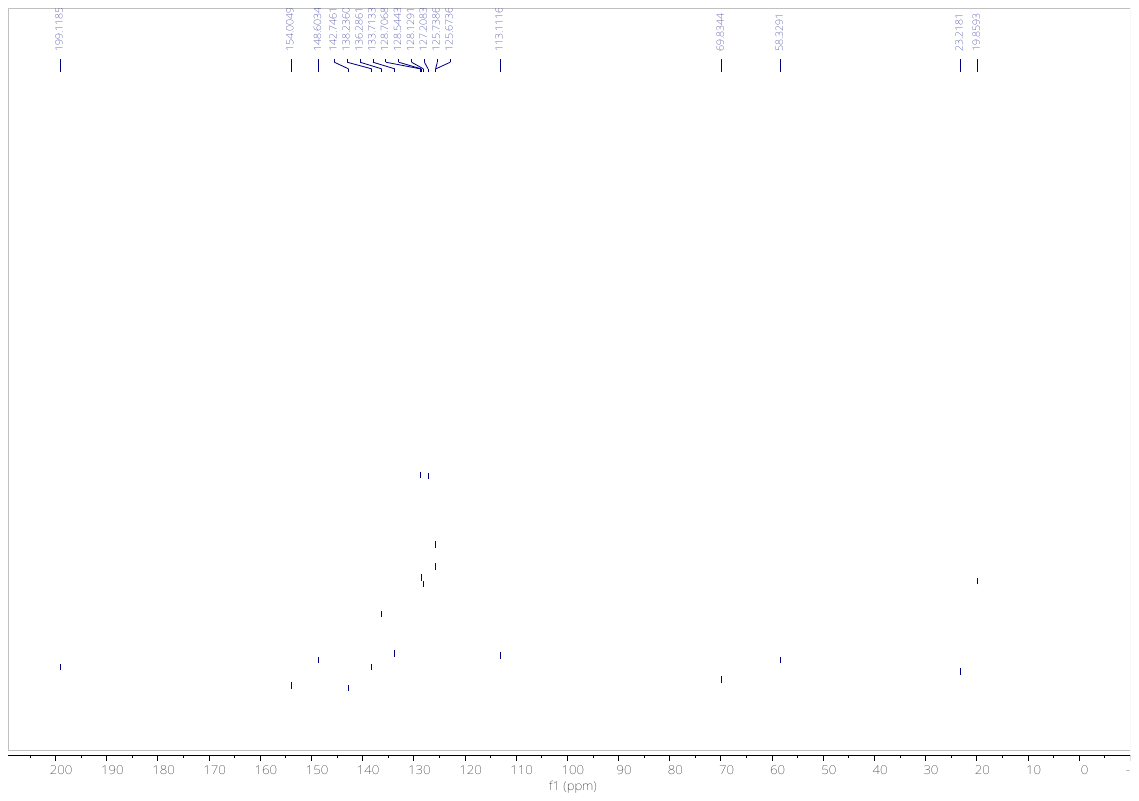

**^13^C NMR** spectrum of **12** (100 MHz, CDCl_3_, 298K)

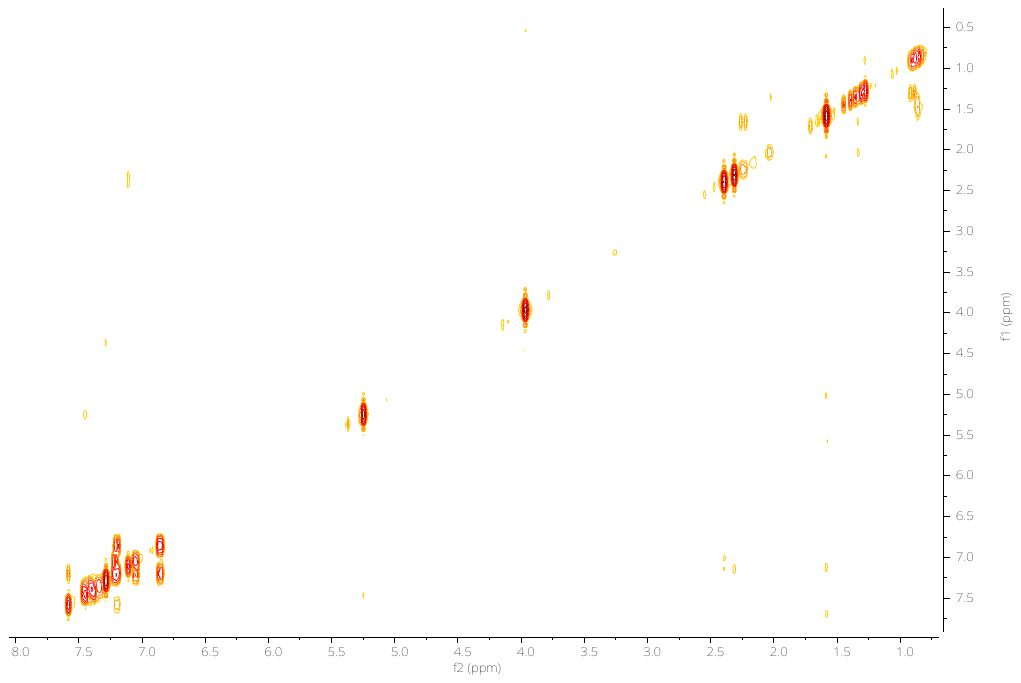

**^1^H-^1^H COSY NMR** spectrum of **12** (400 MHz, CDCl_3_, 298K)

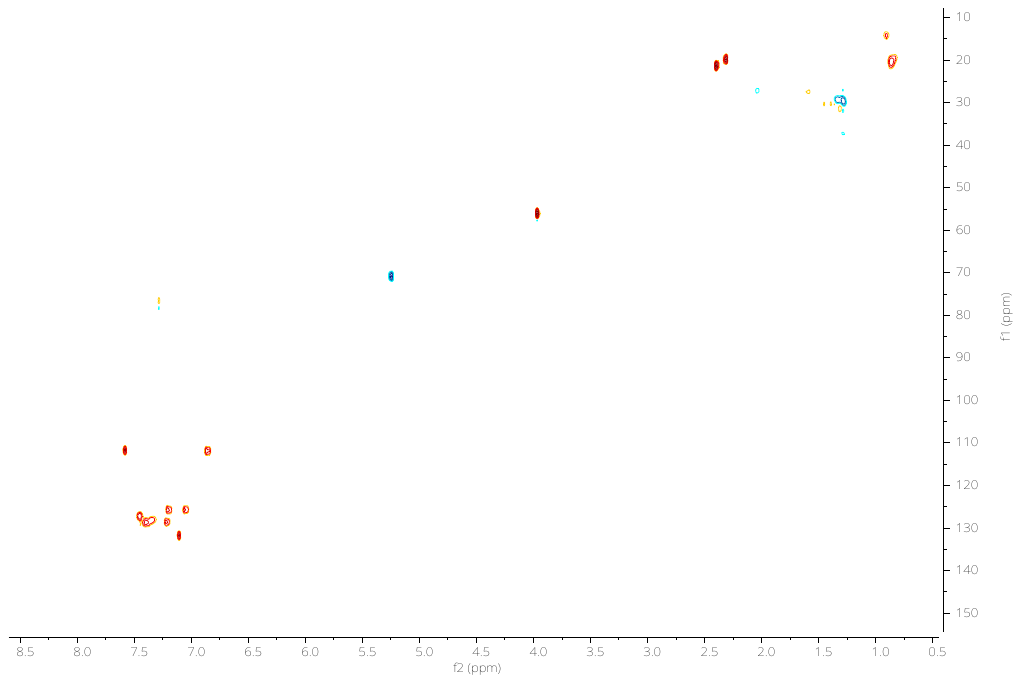

**^1^H-^13^C HSQC NMR** spectrum of **12** (CDCl_3_, 298K)

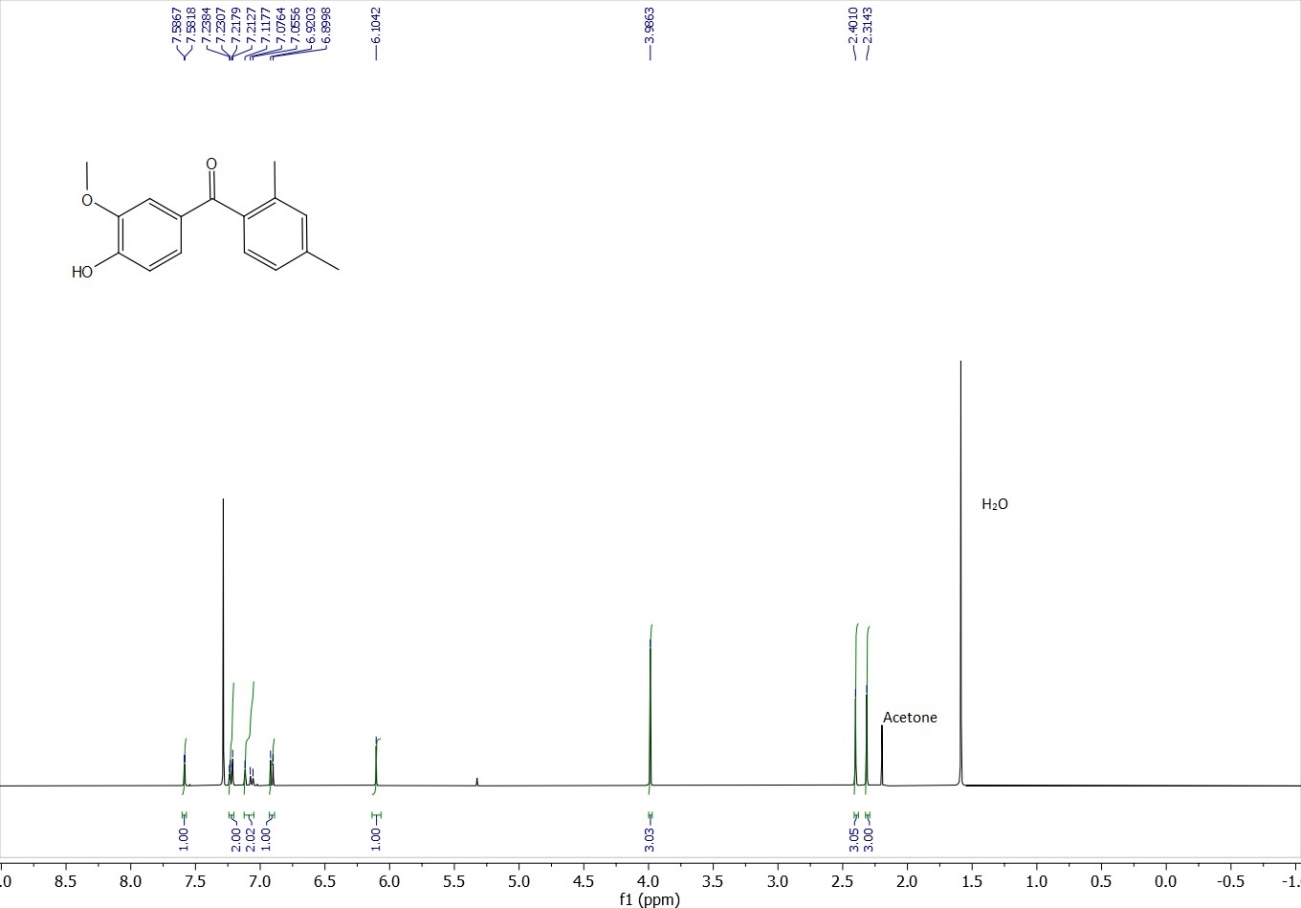

**^1^H NMR** spectrum of **13** (400 MHz, CDCl_3_, 298K)

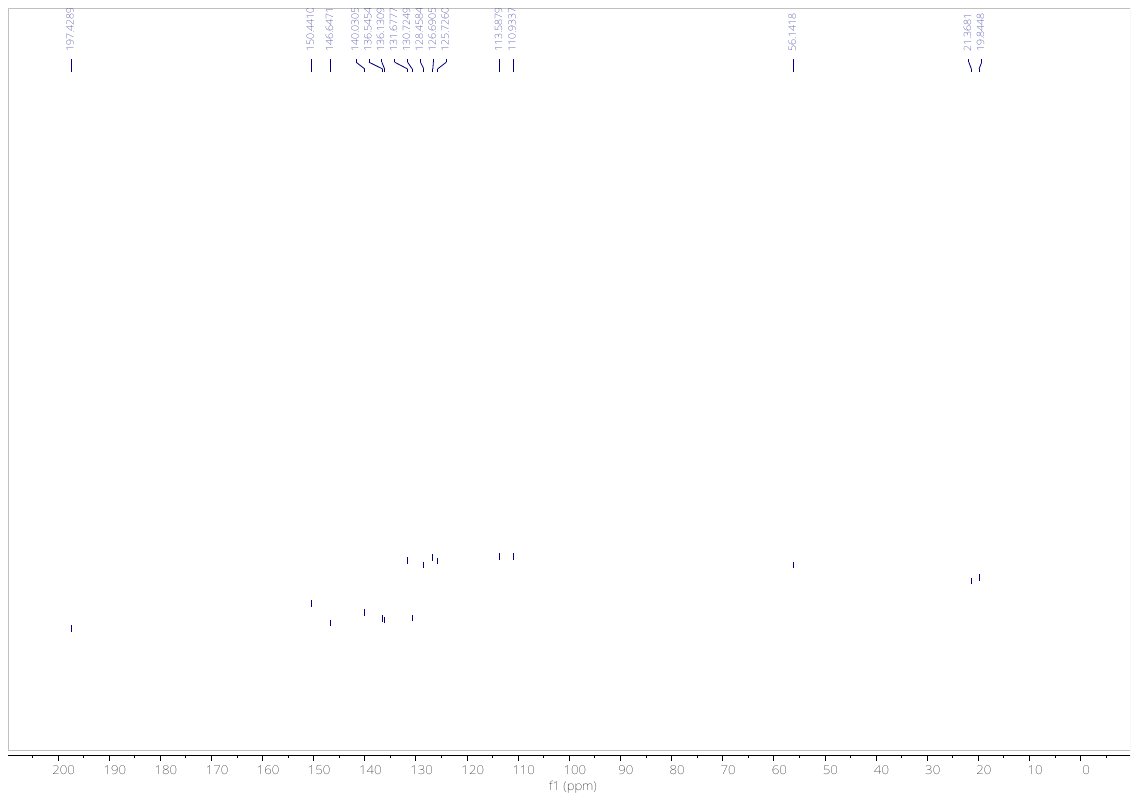

**^13^C NMR** spectrum of **13** (100 MHz, CDCl_3_, 298K)

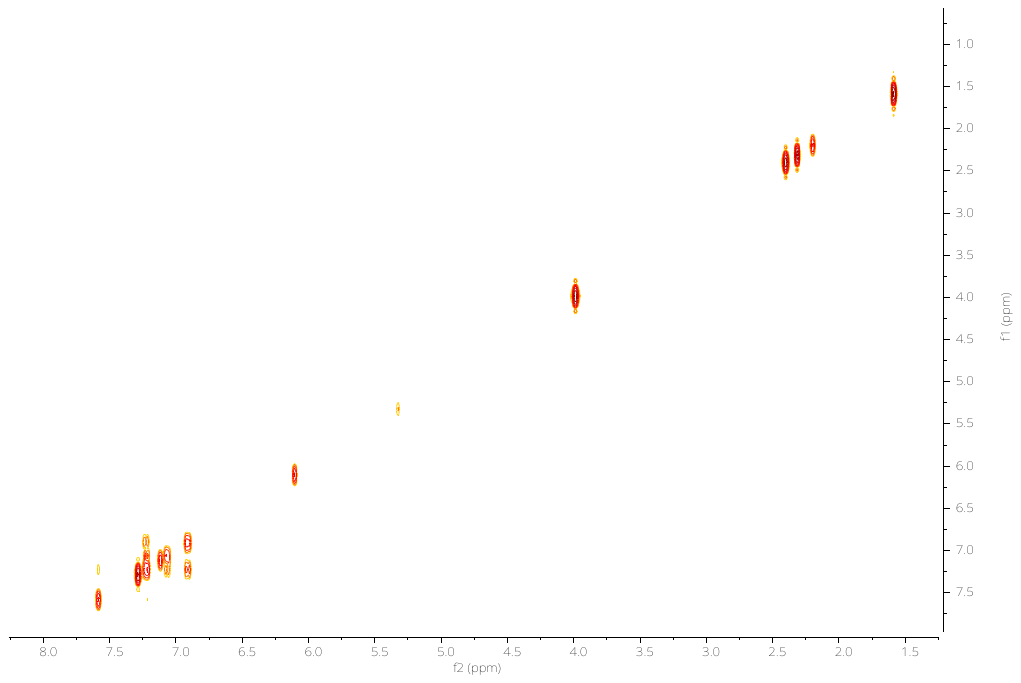

**^1^H-^1^H COSY NMR** spectrum of **13** (400 MHz, CDCl_3_, 298K)

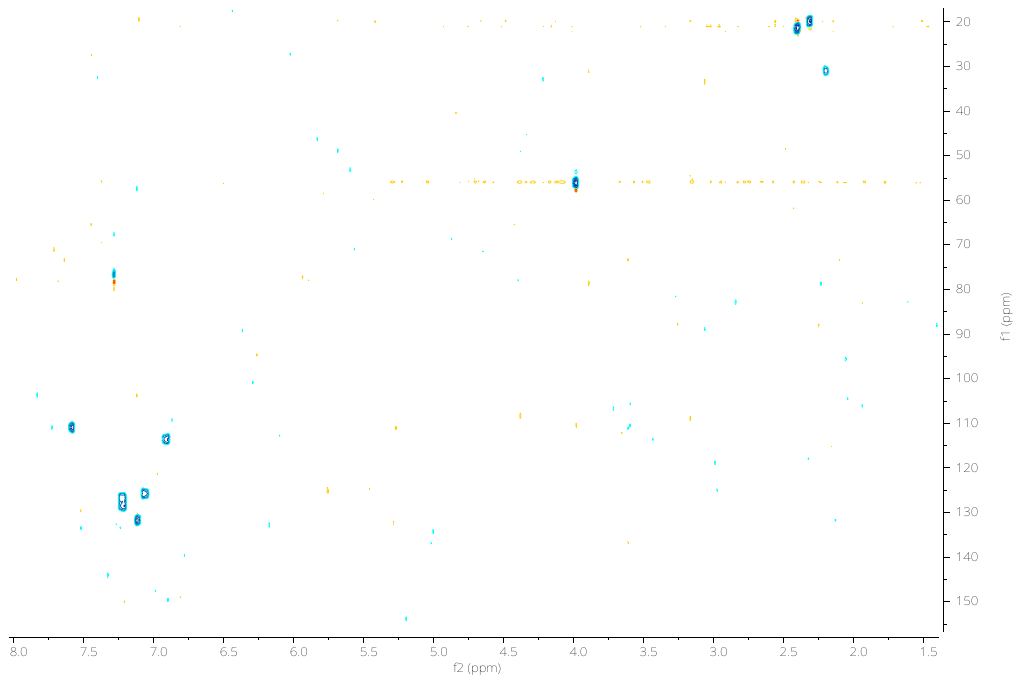

**^1^H-^13^C HSQC NMR** spectrum of **13** (CDCl_3_, 298K)

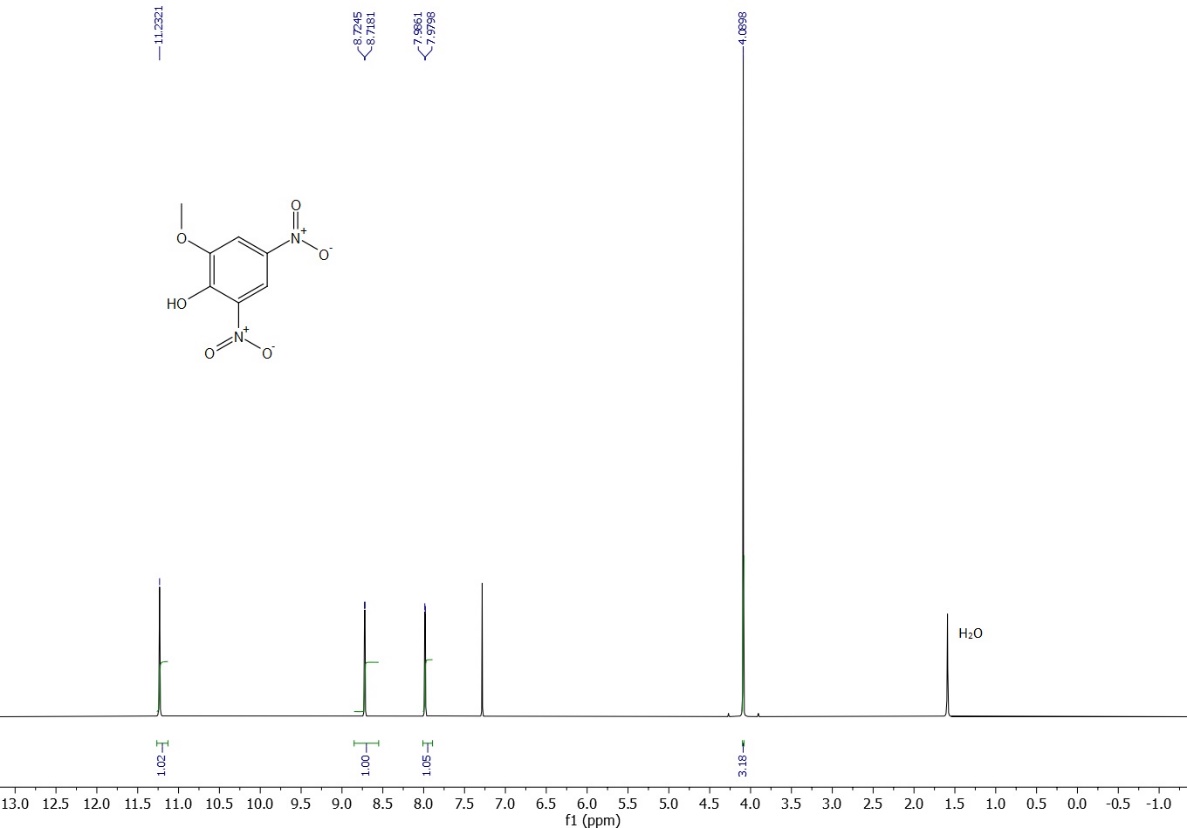

**^1^H NMR** spectrum (400 MHz, CDCl_3_, 298K) of 2,4-dinitro-6-methoxyphenol isolated as product from the nitration of **13**

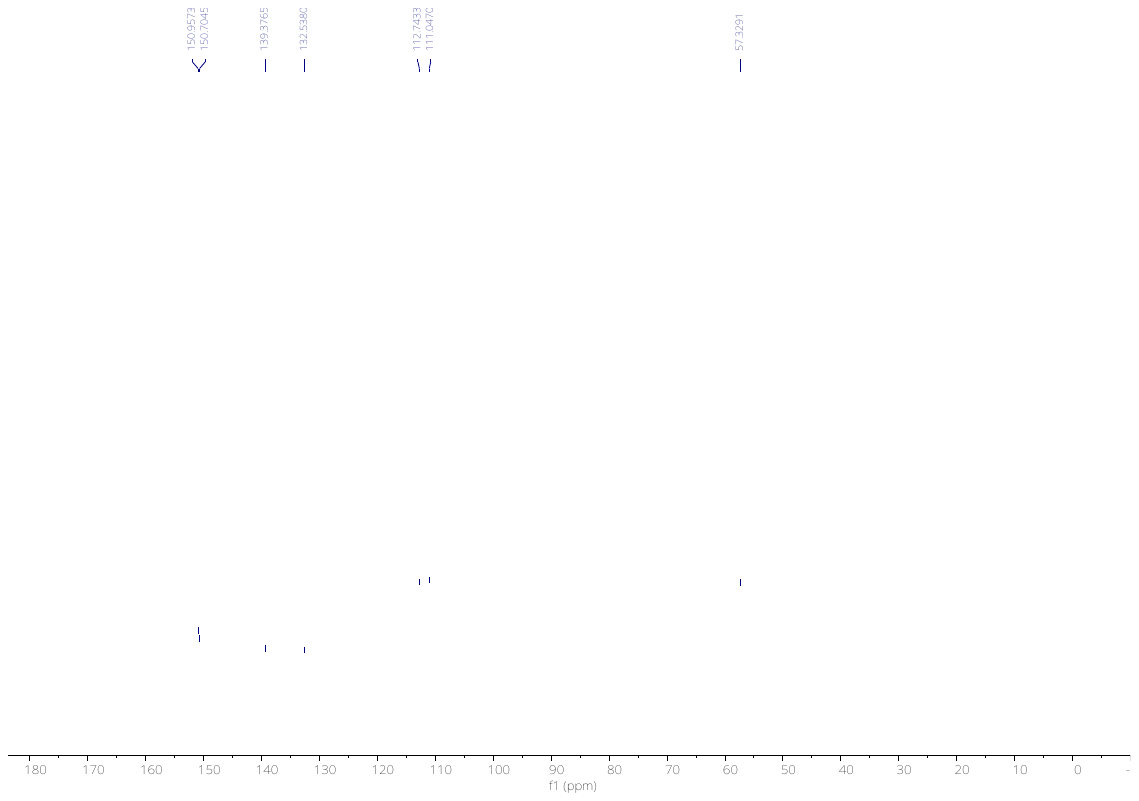

**^13^C NMR** spectrum (100 MHz, CDCl_3_, 298K) of 2,4-dinitro-6-methoxyphenol isolated as product from the nitration of **13**

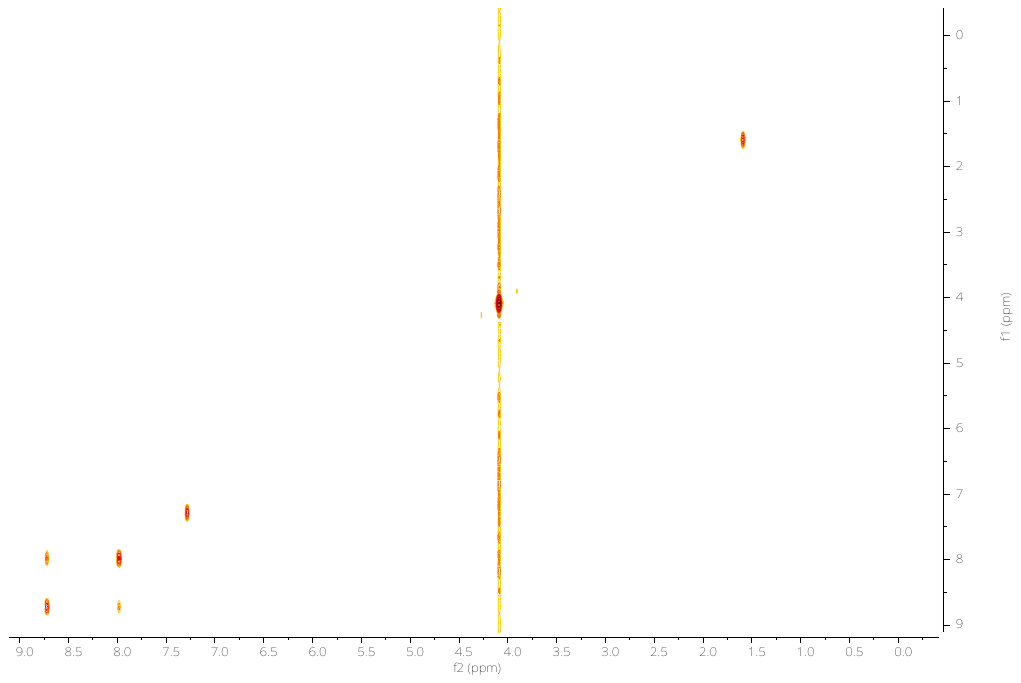

**^1^H-^1^H COSY NMR** spectrum (400 MHz, CDCl_3_, 298K) of 2,4-dinitro-6-methoxyphenol isolated as product from the nitration of **13**

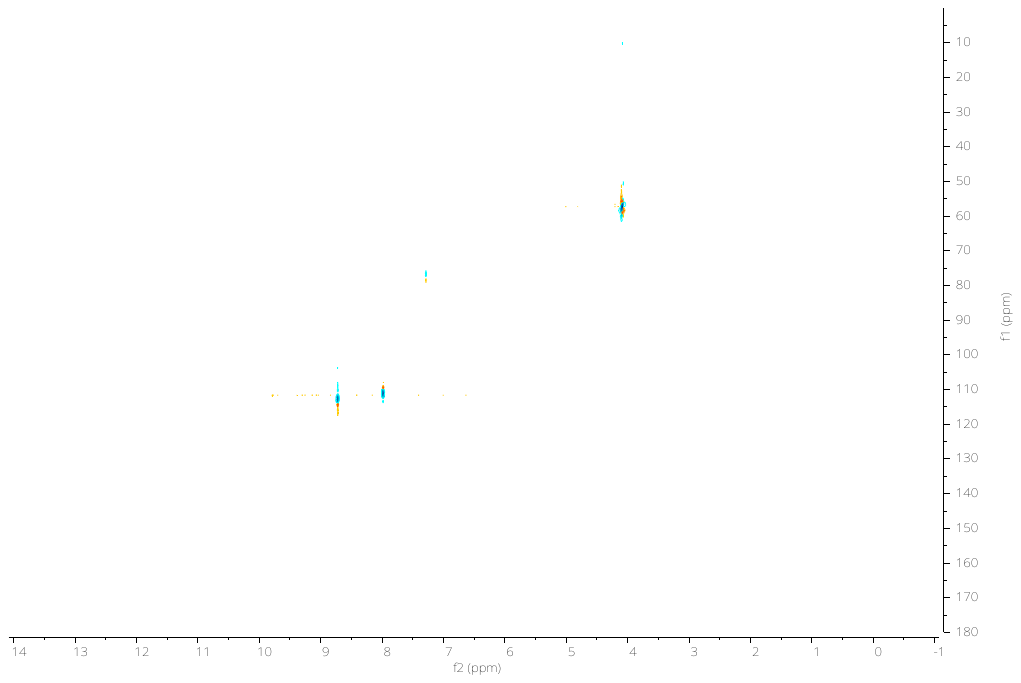

**^1^H-^13^C HSQC NMR** spectrum (CDCl_3_, 298K) of 2,4-dinitro-6-methoxyphenol isolated as product from the nitration of **13**

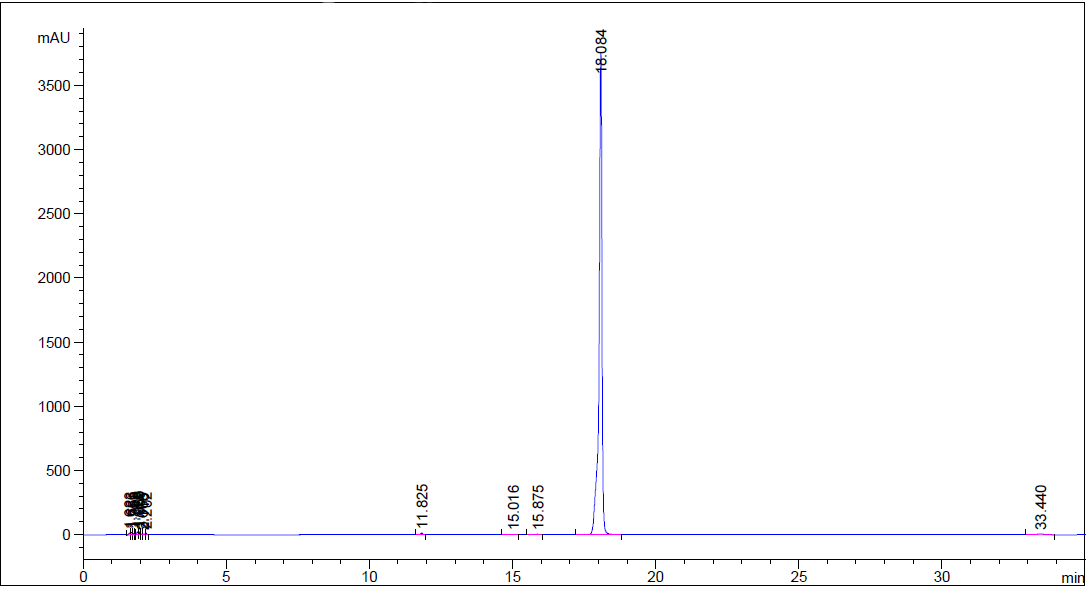

HPLC chromatogram of **1**. The purity degree of compound **1** was determined with an Agilent 1260 HPLC, using a SUPELCO C18 column (3 μm, 150 mm × 4.6 mm) at 40 °C. Mobile phase A: 0.1% TFA in water; mobile phase B: 0.1% TFA in acetonitrile. Gradient conditions: 0−5 min, phase A 100%; 5−15 min, phase A 60%, phase B 40%; 15−25 min, phase A 20%, phase B 80%; 25-30, phase B 100%; 30-35 min, phase A 100%. Flow rate: 1.5 mL/min. The peaks were detected at 270 nm.

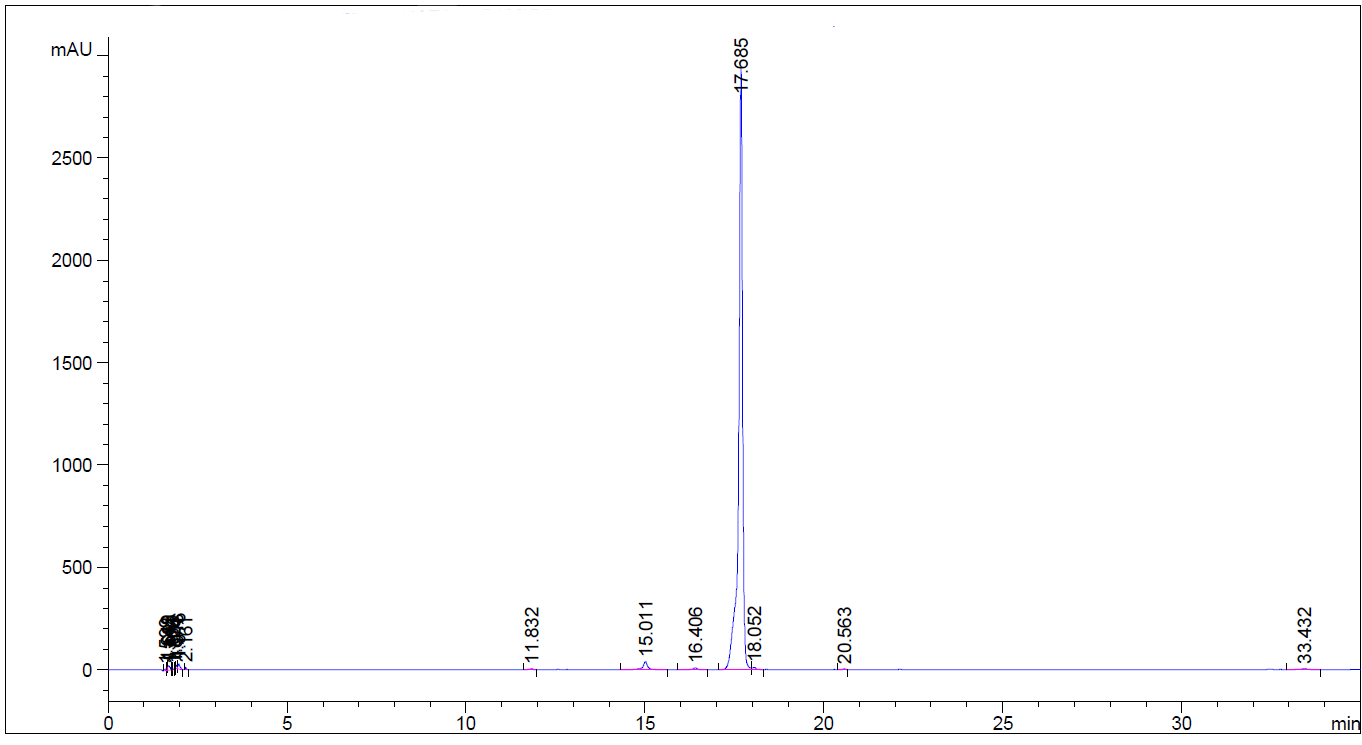

HPLC chromatogram of **2**. The purity degree of compound **2** was determined with an Agilent 1260 HPLC, using a SUPELCO C18 column (3 μm, 150 mm × 4.6 mm) at 40 °C. Mobile phase A: 0.1% TFA in water; mobile phase B: 0.1% TFA in acetonitrile. Gradient conditions: 0−5 min, phase A 100%; 5−15 min, phase A 60%, phase B 40%; 15−25 min, phase A 20%, phase B 80%;25-30, phase B 100%; 30-35 min, phase A 100%. Flow rate: 1.5 mL/min. The peaks were detected at 270 nm.

### **Smiles**

| Compound | SMILES |
| --- | --- |
| **3-OMT** | O=C(C1=CC=C(C)C=C1)C2=CC([N+]([O-])=O)=C(O)C(OC)=C2 |
| **1** | O=C(C1=CC(C)=CC(C)=C1)C2=CC([N+]([O-])=O)=C(O)C(OC)=C2 |
| **2** | O=C(C1=CC=C(C)C=C1C)C2=CC([N+]([O-])=O)=C(O)C(OC)=C2 |

### **Table S2**. Diffraction data collection and refinement statistics.

| **Parameter** | **hTTR / 1** | **hTTR / 2** |
| --- | --- | --- |
| **PDB code** | 8C85 | 8C86 |
| Wavelength (Å) | 0.886 | 0.886 |
| Space group | P 21 21 2 | P 21 21 2 |
| Unit cell (Å) | 42.06, 84.86, 64.28 | 41.96, 84.88, 63.02 |
| Resolution range (Å) | 42.06 - 1.19 (1.21 - 1.19) | 42.44 - 1.10 (1.12 – 1.10) |
| Rmerge | 0.067 (1.205) | 0.082 (1.091) |
| Rpim | 0.028 (0.533) | 0.034 (0.467) |
| Total reflections | 532468 (24570) | 657887 (31379) |
| Unique reflections | 74429 (3634) | 91704 (4420) |
| Multiplicity | 7.1 (6.8) | 7.2 (7.1) |
| Mean((I)/sd(I)) | 11.5 (1.5) | 9.4 (1.6) |
| Mn(I) half-set correlation CC(1/2) | 0.999 (0.520) | 0.995 (0.654) |
| Completeness | 99.9 (100.0) | 99.7 (98.8) |
| Wilson B-factor (Å^2^) | 13.35 | 14.73 |
| Reflections used in refinement | 74418 (7367) | 91691 (8956) |
| Reflections used in free set | 3746 (352) | 4648 (423) |
| R_work_ | 0.142 (0.254) | 0.1381 (0.2378) |
| R_free_ | 0.168 (0.288) | 0.1581 (0.2677) |
| RMSD bonds (Å) | 0.019 | 0.014 |
| RMSD angles (°) | 1.62 | 1.30 |
| Ramachandran favoured (%) | 97.4 | 97.8 |
| Ramachandran allowed (%) | 2.6 | 2.2 |
| Ramachandran outliers (%) | 0 | 0 |
| Rotamer outliers | 2.4 | 0.88 |
| Overall number of atoms (non-H) | 2307 | 2176 |
| in macromolecules | 2078 | 1962 |
| in ligands | 70 | 74 |
| in solvent | 185 | 170 |
| Average B-factor (Å^2^) | 19.22 | 21.94 |
| for macromolecules | 17.85 | 20.84 |
| for ligands | 20.92 | 20.68 |
| for solvent | 34.17 | 34.87 |
